# A nuclear CobW/WW-domain factor represses the CO_2_-concentrating mechanism in the green alga *Chlamydomonas reinhardtii*

**DOI:** 10.1101/2025.07.09.663896

**Authors:** Daisuke Shimamura, Junko Yasuda, Yosuke Yamahara, Hirofumi Nakano, Shin-Ichiro Ozawa, Ryutaro Tokutsu, Ayumi Yamagami, Tomonao Matsushita, Yuichiro Takahashi, Takeshi Nakano, Hideya Fukuzawa, Takashi Yamano

## Abstract

Microalgae induce a CO_2_-concentrating mechanism (CCM) to maintain photosynthesis when dissolved CO_2_ is limited, but how this energy-intensive system is suppressed when CO_2_ levels rise has remained unclear. The CCM consumes 15–30% of photosynthetically-generated ATP, making its regulation critical for cellular energy balance. Here, we identify a nuclear repressor of the CCM in the green alga *Chlamydomonas reinhardtii*. A pull-down screen for interacting partners of the master activator CCM1/CIA5 revealed an uncharacterized protein that co-purifies with CCM1 even after high-salt washes. This protein, designated CCM1-binding protein 1 (CBP1), combines a CobW/CobW_C GTP-binding metallochaperone module with a WW domain characteristic of protein-protein interactions. CBP1 colocalizes with CCM1 in the nucleus regardless of CO_2_ conditions, and the two proteins interact *in vivo*. CRISPR/Cas9-based disruption of CBP1 does not affect growth or CCM induction under CO_2_ limitation but derepresses 27 of 41 CCM1-dependent low-CO_2_ inducible genes under high-CO_2_ conditions. These include the periplasmic and intracellular carbonic anhydrases (CAH1, LCIB) and inorganic carbon transporters/channels (LCIA, LCI1, BST1, BST3). Consistently, *cbp1* mutants accumulate higher levels of CAH1 and LCIB proteins and exhibit a 40% increase in inorganic carbon affinity under high-CO_2_ conditions; this phenotype is rescued by CBP1 complementation or by acetazolamide treatment. These results demonstrate that CBP1 prevents unnecessary CCM activity when CO_2_ is abundant, acting upstream of both transporter/channel and carbonic anhydrase modules. Our findings suggest a regulatory mechanism potentially linking zinc-dependent protein chemistry to CCM gene repression, providing insights into energy-efficient CO_2_ sensing in aquatic photosynthetic organisms.

**Significance statement:** Algae flourish by activating a CO_2_-concentrating mechanism (CCM) when dissolved CO₂ is limited but must deactivate it when CO_2_ levels increase to conserve energy. We have identified the nuclear protein that functions as this long sought “off switch” in the model green alga. Deletion of this protein causes cells to overproduce CCM transporters and enzymes, maintaining CCM activity even under high CO_2_ conditions, demonstrating its essential role in suppressing the system when carbon is abundant. This finding illuminates how algae balance energy consumption with carbon capture and offers a new target for engineering strains that fix CO_2_ more efficiently for biofuel production or climate-mitigation technologies.

## Introduction

Efficient CO_2_ sensing enables photosynthetic organisms to maximize carbon acquisition while minimizing resource expenditure. In terrestrial plants, stomata regulate their aperture in response to environmental CO_2_, balancing photosynthesis with transpiration (1). A recently identified guard-cell complex comprising the Raf-like kinase HT1 and MAP kinases MPK12/MPK4 now provides a molecular model for CO_2_/HCO_3_^−^ sensing (2). These discoveries highlight the evolutionary diversity of CO_2_-responsive pathways.

Aquatic photoautotrophs encounter chronic inorganic carbon (Ci) limitation because dissolved CO_2_ rapidly converts to HCO_3_^−^. Many microalgae, including marine diatoms and freshwater algae, therefore employ a CO_2_-concentrating mechanism (CCM) that utilizes Ci transporters and carbonic anhydrases (CAs) to accumulate CO_2_ around the photosynthetic CO_2_-fixing enzyme, Ribulose-1,5-bisphosphate carboxylase/oxygenase (Rubisco), enhancing CO_2_ fixation (3, 4). Despite being the primary carbon fixation enzyme, Rubisco has relatively low affinity and specificity for CO_2_, making CCM essential under CO_2_-limiting conditions (5). The green alga *Chlamydomonas reinhardtii* (hereafter *Chlamydomonas*) strongly induces Ci transporters, CAs, and regulatory proteins such as low-CO_2_ response regulator 1 (LCR1) when CO_2_ becomes limiting (6, 7).

Central to CCM regulation in *Chlamydomonas* is CCM1/CIA5 (hereafter CCM1), a zinc-binding protein identified independently by two research groups (8, 9). Loss-of-function mutants of CCM1 fail to induce essential CCM genes under CO_2_-limiting conditions (6), resulting in severe growth defects. Genome-wide analyses further reveal that many CCM1-dependent genes respond to additional factors beyond CO_2_ concentration, such as light intensity, photoperiod, and nitrogen availability (10–12). Although CCM1 contains two Zn^2+^-binding sites (13, 14), direct DNA binding has not been demonstrated, suggesting that CCM1 operates in conjunction with other nuclear factors.

In contrast, mechanisms that deactivate the CCM when CO_2_ again becomes abundant remain poorly characterized. In cyanobacteria, the LysR regulators NdhR/CcmR and CmpR repress Ci-transporter operons under high CO_2_ (15, 16). A CREB/ATF-type bZIP factor performs a similar function in the diatom *Phaeodactylum tricornutum* (17). In green algae, several mutants with impaired CCM induction have been described, yet no dedicated nuclear “off-switch” has been confirmed. This gap is significant because CCM operation can consume approximately 15–30% of photosynthetically generated ATP, as demonstrated by direct energy-budget measurements (18) and consistent with modeling of pyrenoid-based CCM energetics (19), indicating a substantial energetic trade-off between CCM operation and other cellular processes.

Metal-dependent transcriptional switches represent a plausible mechanism for such control. Members of the COG0523 GTPase family, including plant ZNG1/2, deliver Zn^2+^ to target proteins and modulate gene expression according to zinc status (20–22). Similar Zn-based sensors regulate diverse stress pathways in higher plants—for example, the bZIP19/23 system that activates ZIP4 under Zn deficiency (23). Since CCM1 itself coordinates Zn^2+^ (13), it is conceivable that changes in intracellular Zn availability could rapidly trigger a conformational change in CCM1, modulating its interaction with regulatory partners and thus providing a quick molecular switch under high CO_2_ conditions, similar to known Zn-regulated transcription factors (24).

In this study, we tested the hypothesis that a nuclear factor binds CCM1 and inhibits CCM gene expression under high CO_2_. A pull-down screen indeed identified a CobW/WW-domain protein that we name CCM1-binding protein 1 (CBP1). We demonstrate that CBP1 associates with CCM1 *in vivo*, localizes to the nucleus, and is specifically required to repress a subset of CCM genes when CO_2_ is abundant. Our findings reveal a previously unidentified layer of CCM regulation that links zinc metabolism to energy conservation in aquatic phototrophs, addressing a regulatory aspect previously overlooked in algal CCM studies.

## Results

### Identification of the CCM1-binding proteins

To identify proteins that physically interact with the master regulator CCM1, we performed a pull-down assay using *Chlamydomonas reinhardtii* cells expressing FLAG-tagged CCM1. Specifically, we generated a transgenic strain (CF-2) by introducing a CCM1-FLAG construct into the *ccm1* mutant strain C16 (8) (Fig. S1A). We chose heterotrophic conditions for cell growth to minimize background expression of CCM-related genes and to stabilize CCM1 expression for immunoprecipitation. We then compared immunoprecipitates obtained from the parental strain (5D) and CF-2.

Pull-down experiments revealed that the CCM1-FLAG complex contained two major components beyond CCM1 itself (Fig. 1A). In CF-2 cells, two distinct protein bands of approximately 75 kDa and 45 kDa were visible on SDS-PAGE. LC-MS/MS identified the 75 kDa band as a previously uncharacterized protein encoded by *Cre16.g684650*, while the 45 kDa band corresponded to glutamine dehydrogenase 2 (GDH2) (Fig. S2 and Table S1 and S2). To assess binding strength, we performed washes with increasing KCl concentrations (0.1, 0.5, and 1.5 M). A 1.5 M KCl wash abolished the GDH2 signal but not the 75 kDa protein, indicating that GDH2 associates only weakly with CCM1, whereas the 75 kDa protein forms a tight complex (Fig. 1A). We therefore named the 75 kDa protein as CCM1-binding protein 1 (CBP1). A parallel analysis using in-solution tryptic digestion of the entire CCM1-FLAG eluate corroborated these findings, confirming the presence of CBP1 and GDH2, and additionally identifying glutamate dehydrogenase 1 (GDH1) as another component of the complex (Fig. S2 and Table S3).

**Fig. 1.**
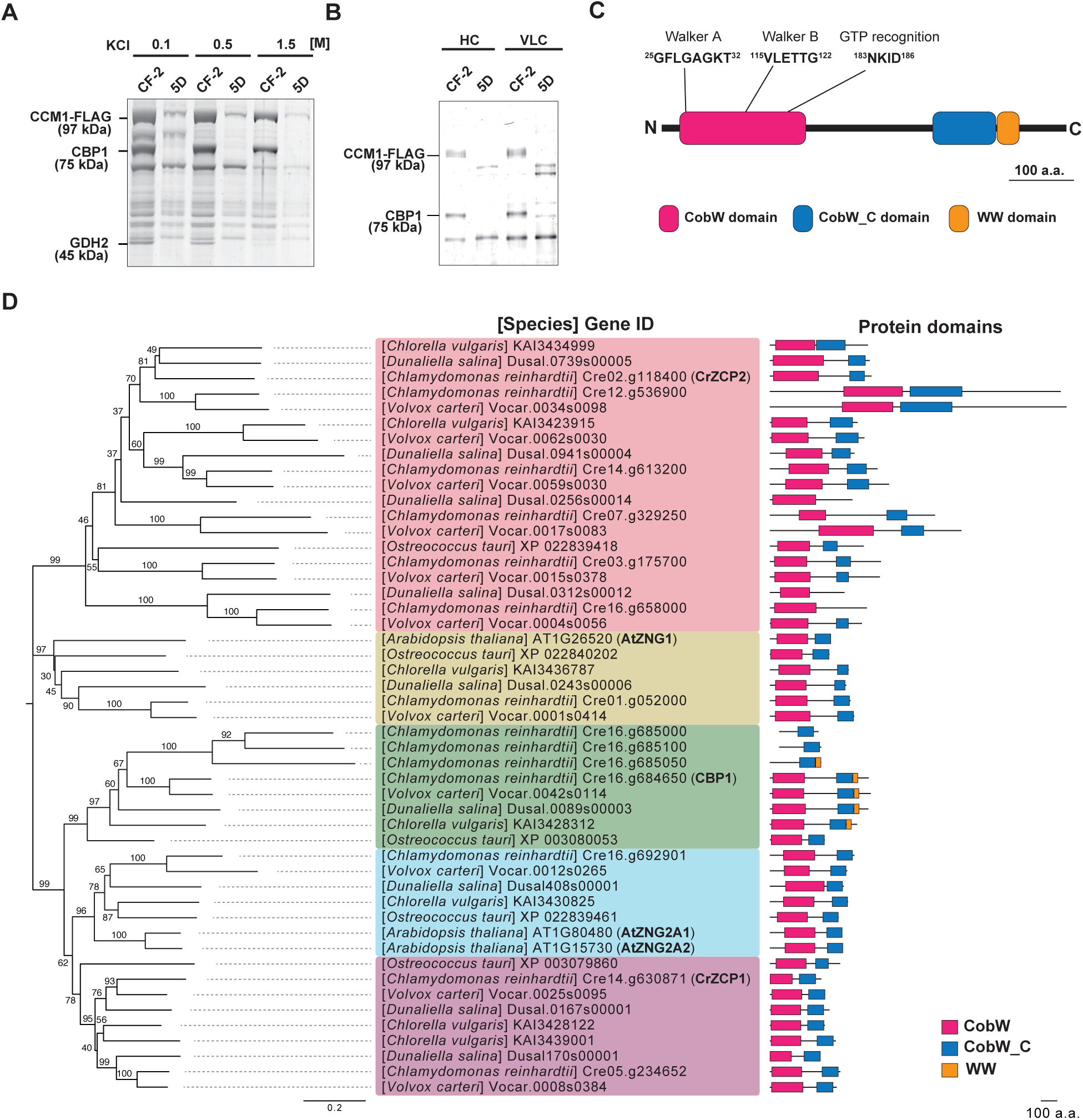
CBP1, a CobW/WW domain-containing protein, is identified as a CCM1-interacting partner. (A) The CCM1-FLAG protein complex was isolated by pull-down assay and subjected to SDS-PAGE followed by silver staining. The arrows indicate the positions of the major components of the complex, with their estimated molecular weights. Cells were cultured under heterotrophic conditions. The KCl concentration in the wash buffer is indicated. (B) Accumulation of CBP1 in CCM1-FLAG complexes isolated from C16:CCM1-FLAG line 2 (CF-2) cells grown under HC (5% CO_2_) and VLC (0.04% CO_2_) conditions. 5D represents the parental strain used to generate CF-2. CBP1 was detected by immunoblot analysis using anti-CBP1 antibody. CCM1-FLAG was detected using anti-FLAG antibody. (C) Domain structure and motif sequence of the CBP1 protein. The CobW domain contains conserved amino acid motifs important for GTPase activity including the P-loop (GxxxxGKT), Switch I (DxxG), and Switch II (NKxD) motifs. (D) Phylogenetic tree of COG0523 proteins conserved in *Arabidopsis thaliana*, *Chlamydomonas reinhardtii*, *Volvox carteri*, *Dunaliella salina*, *Chlorella vulgaris* and *Ostreococcus tauri*. The phylogenetic tree was constructed using the Neighbor-Joining (NJ) method based on the amino acid sequences of CBP1 homologs. Bootstrap values (1,000 replicates) are shown at the nodes. The scale bar of the phylogenetic tree represents 0.2 substitutions per site. Protein domains are illustrated to the right of each sequence: CobW (pink), CobW_C (blue), and WW (orange) domains. CBP1 and ZNG1 orthologs are highlighted in bold.

Subsequent immunoblot analyses showed that CBP1 is present at similar levels in cells grown under both HC and very-low-CO_2_ (VLC) conditions (Fig. 1B). Analysis of the immunoprecipitated CCM1 complex also revealed several potential phosphorylation sites on CCM1 itself (Table S4). Furthermore, a reciprocal co-immunoprecipitation assay demonstrated that an anti-CBP1 antibody successfully co-precipitated CCM1 under both HC and VLC conditions (Fig. S1B). Together, these results suggest that the CCM1-CBP1 interaction occurs constitutively and does not depend on external CO_2_ levels. These findings prompted us to investigate CBP1’s role in CCM regulation under varying CO_2_ conditions.

### CBP1 contains a CobW/CobW_C domain and a WW domain

Sequence analysis revealed that CBP1 possesses three recognizable domains: a CobW domain, a CobW_C domain, and a WW domain (Fig. 1C). The CobW domain contains conserved Walker A (residues 25-32: GFLGAGKT) and Walker B (residues 115-122: VLETTG) motifs, as well as a GTP recognition sequence (residues 183-186: NKID), all essential for GTPase activity. Proteins harboring CobW/CobW_C typically belong to the COG0523 family, known for metal chaperone activity (20). In Viridiplantae, two COG0523-type proteins—Zn-regulated GTPase metalloprotein activator 1 (ZNG1) and ZNG2—are broadly conserved (22). In *Chlamydomonas*, ZCP1 and ZCP2 similarly encode COG0523 proteins that are induced under zinc-deficient conditions (20). Unlike these characterized COG0523 proteins, CBP1 also features a C-terminal WW domain, which typically mediates protein-protein interactions through proline-rich sequences.

Phylogenetic analysis of COG0523 proteins across green algae and plants indicates that CBP1 belongs to a separate clade from ZNG1/2, and that WW-domain-containing COG0523 proteins are restricted to certain green algal lineages (Chlorophyceae and Trebouxiophyceae), precisely mirroring CCM1’s taxonomic distribution (Fig. 1D and Fig. S3). This co-occurrence strongly suggests co-evolution of the CBP1-CCM1 regulatory module. These observations suggest that CBP1 exerts a distinct nuclear function in *Chlamydomonas*, potentially involving metal ion transfer and protein-binding events.

To test CBP1’s role in CCM regulation, we generated a *cbp1* mutant by CRISPR/Cas9-directed insertion of an *AphVII* resistance cassette into the second exon of *CBP1* in the wild-type strain C9 (Fig. 2A). Genomic PCR confirmed the successful insertion of the cassette, with the mutant showing a 3.0 kb band compared to the 1.0 kb wild-type band (Fig. 2B).

**Fig. 2.**
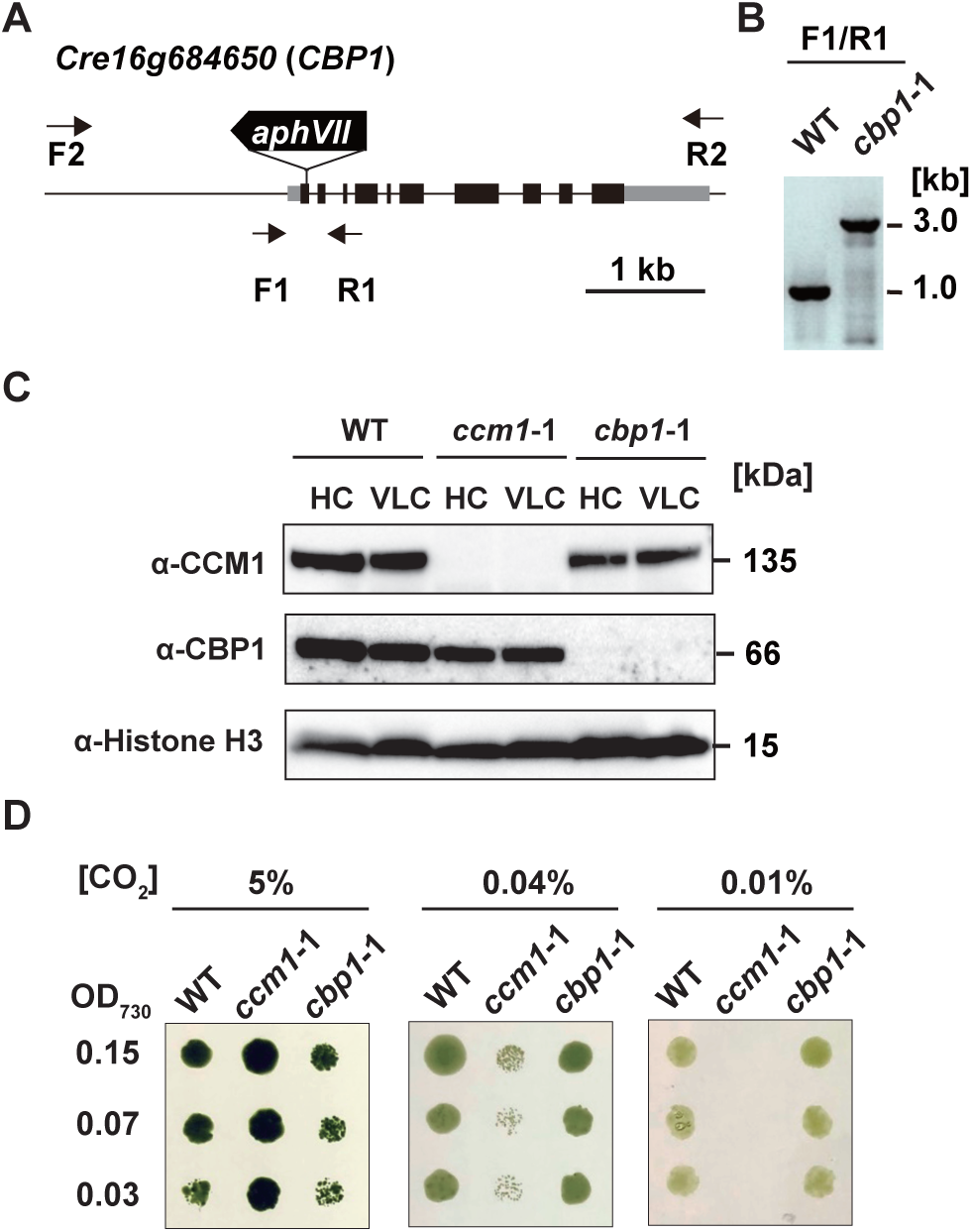
Loss of CBP1 does not affect growth under various CO_2_ conditions. (A) Schematic representation of the *aphVII* cassette insertion in the *cbp1* mutant. Exons, introns, and untranslated regions are shown as black boxes, black lines, and gray boxes, respectively. The *aphVII* cassette conferring paromomycin resistance is shown as a white box with an arrow indicating its orientation. Binding sites of PCR primers are indicated as arrows. (B) Genomic PCR to confirm insertion of the *aphVII* cassette in the *CBP1* gene. (C) Accumulation of CCM1 and CBP1 in WT, *ccm1* mutant (*ccm1*-1), and *cbp1* mutant (*cbp1*-1). Cells grown in 5% CO_2_ were transferred to 5% (HC) or 0.04% (VLC) CO_2_ for 12 h. Histone H3 was used as a loading control. (D) Spot tests of WT, *ccm1*-1, and *cbp1*-1. Cells grown to logarithmic phase were diluted to the indicated optical density (OD_730_ = 0.30, 0.15, or 0.07), and the cell suspensions were spotted on MOPS-P agar plates. Cells were grown in closed chambers supplied with 5%, 0.04%, or 0.01% CO_2_ under continuous light at 120 μmol photons m^−2^ s^−1^. Images were taken after 3 days of growth.

To examine protein accumulation, we analyzed CCM1 and CBP1 levels under high CO_2_ (HC, 5%) and very-low CO_2_ (VLC, 0.04%) conditions. Immunoblotting with an anti-CBP1 antibody confirmed that the resulting mutant (*cbp1*-1) completely lacked CBP1 protein under both HC and VLC conditions (Fig. 2C). Importantly, CCM1 protein levels in the *cbp1*-1 mutant were comparable to those in wild-type under VLC conditions. Likewise, CBP1 protein levels in the *ccm1*-1 mutant (25) were also similar to those in wild-type under both HC and VLC conditions (Fig. 2C). These results demonstrate that the accumulation (or stability) of CCM1 and CBP1 does not require their interaction.

Growth analysis revealed striking differences between the *cbp1* and *ccm1* mutants. While *ccm1*-1 cells showed severe growth defects at 0.04% and 0.01% CO_2_, *cbp1*-1 cells grew normally at all CO_2_ concentrations tested (5%, 0.04%, and 0.01%) (Fig. 2D). The potential implications of the subtle growth differences observed under HC conditions are considered further in the Discussion. This normal growth phenotype indicates that CBP1, unlike CCM1, is dispensable for CCM induction and cell survival under low CO_2_ conditions. Thus, CBP1 is dispensable for growth under both high and low CO_2_ conditions, implying a function distinct from that of CCM1.

Attempts to create a CRISPR/Cas9-directed *GDH1/2* insertion mutants with four independent guide RNAs failed to yield viable transformants (> 1,000 colonies screened), suggesting GDH1/2 is essential under our culture conditions and may relay metabolic information to CCM1; this possibility is considered further in the Discussion.

### CBP1 interacts with CCM1 in the nucleus

Because CCM1 localizes to the nucleus, we hypothesized that CBP1 would function in the same compartment. To visualize CBP1’s localization, we fused CBP1 to the fluorescent protein mGold and introduced the construct into *cbp1*-1, generating *cbp1*-1:*CBP1*-mGold. In parallel, we fused CCM1 to mGold and introduced it into *ccm1*-1, yielding *ccm1*-1:*CCM1*-mGold. Confocal microscopy revealed strong nuclear signals in both transformants under HC and VLC conditions (Fig. 3A). The mGold signals clearly localized to a distinct nuclear region that did not overlap with chlorophyll autofluorescence, as shown in the merged images. Notably, the nuclear localization pattern remained unchanged between HC and VLC conditions for both proteins, with the enlarged images showing punctate nuclear signals characteristic of transcription factor localization (Fig. 3A).

**Fig. 3.**
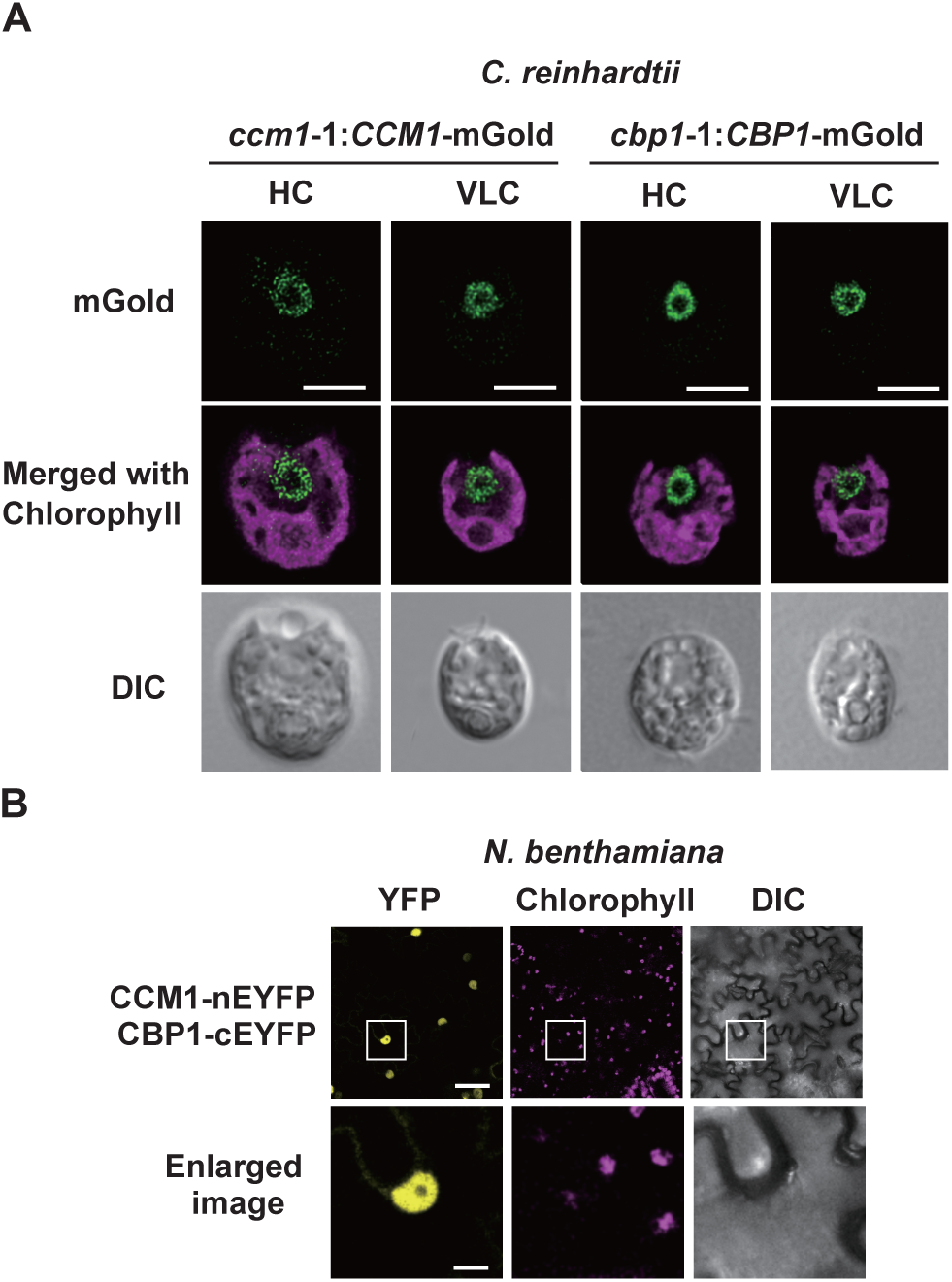
CBP1 localizes to the nucleus and physically interacts with CCM1. (A) Intracellular localization of CCM1 and CBP1-mGold fusion proteins in *Chlamydomonas reinhardtii* cells. Cells were observed under conditions acclimated to either 5% (HC) or 0.04% CO_2_ (VLC) for 12 h. Yellow fluorescence indicates the localization of fusion proteins. Chl represents chlorophyll autofluorescence. DIC shows differential interference contrast images. Scale bars represent 5 μm. (B) BiFC assay to evaluate the nuclear interaction between CCM1 and CBP1. Co-expression of CCM1-nYFP and CBP1-cYFP in leaf epidermal cells of *Nicotiana benthamiana* was analyzed 3 days after infiltration. Reconstituted YFP fluorescence indicates protein-protein interaction. Nuclear localization was confirmed by DAPI staining. Merge shows overlay of YFP and DAPI signals. Scale bars indicate 50 µm (upper panel) and 10 μm (bottom panel), respectively.

To confirm *in vivo* CCM1-CBP1 interactions, we employed bimolecular fluorescence complementation (BiFC), in which YFP is split into nEYFP and cEYFP fragments that fluoresce only when brought together by an interacting protein pair. We tagged CCM1 and CBP1 with nEYFP and cEYFP, respectively, and transiently expressed them in *Nicotiana benthamiana* leaves. A bright nuclear YFP signal was observed exclusively when CCM1-nEYFP and CBP1-cEYFP were co-expressed (Fig. 3B). Together with our pull-down results, these data demonstrate that CCM1 and CBP1 physically interact in the nucleus, regardless of external CO_2_ availability.

### CBP1 represses several CCM-related genes under HC conditions

To determine whether CBP1 influences CCM gene expression, we performed RNA-seq on strains lacking or complementing both CCM1 and CBP1 under HC and VLC. Specifically, we generated *ccm1*-1:*CCM1* by introducing a functional *CCM1* genomic fragment into *ccm1*-1, selecting a transformant whose Ci-affinity under VLC matched that of WT (Table S5). Likewise, *cbp1*-1:*CBP1* was established by reintroducing *CBP1* genomic fragment into *cbp1*-1.

We then cultured WT, *ccm1*-1, *cbp1*-1, *ccm1*-1:*CCM1*, and *cbp1*-1:*CBP1* under HC for 24 h, shifted them to VLC for 0.3 or 2.0 h, and analyzed their transcriptomes using Trimmed Mean of M-values (TMM) normalization. We first defined VLC-inducible genes as those significantly upregulated (FDR < 0.01, log_2_ fold change > 1) after 0.3 or 2.0 h under VLC in WT. A principal component analysis (PCA) of these VLC-inducible genes revealed that *cbp1*-1 displayed a distinct expression profile compared to WT or *ccm1*-1, with HC samples clustering separately from VLC samples along PC1 (62.6% variance), while *cbp1*-1 samples under HC diverged from other HC samples along PC2 (16.4% variance), pointing to a unique role for CBP1 (Fig. 4A).

**Fig. 4.**
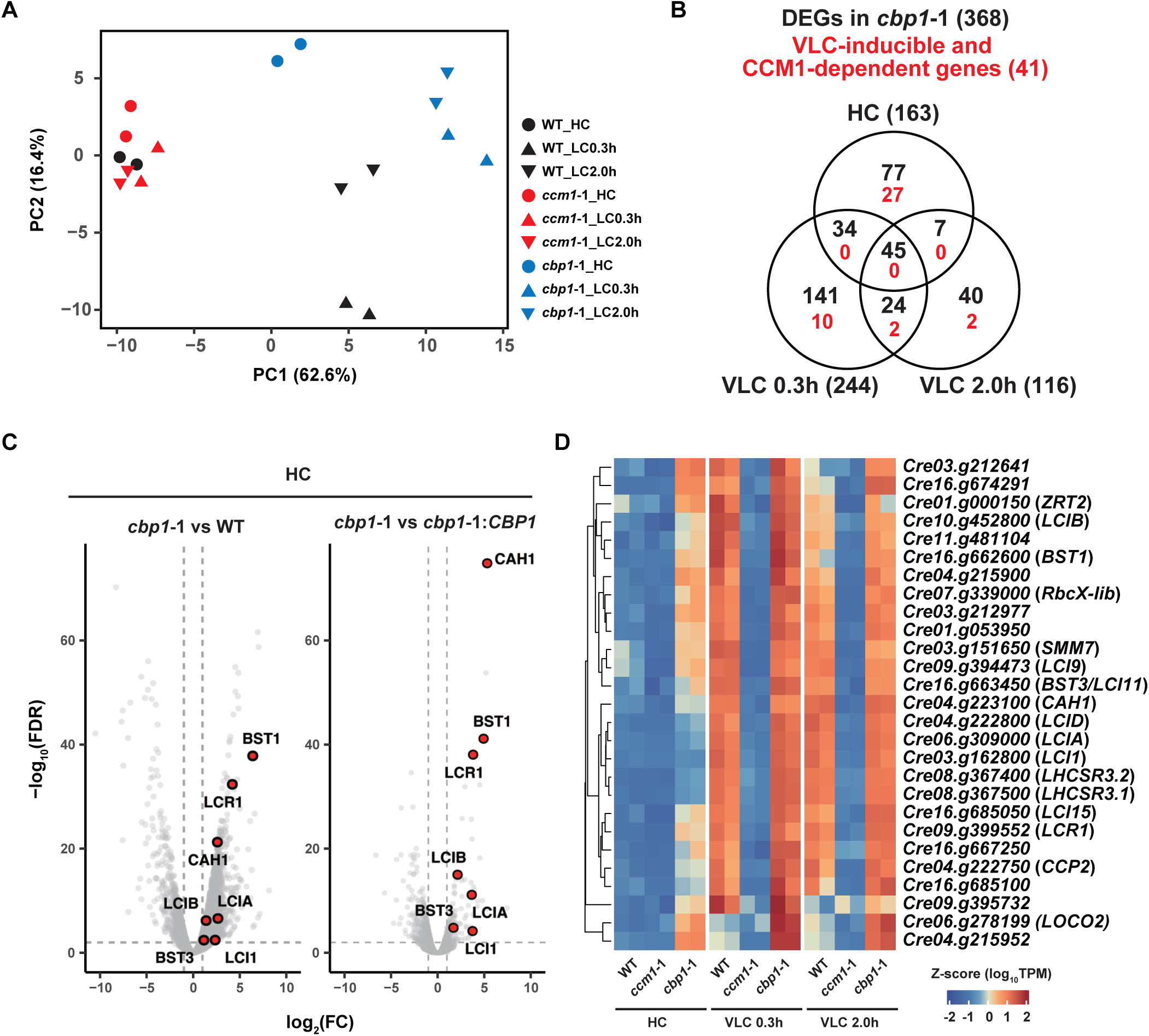
CBP1 represses CCM gene expression under high CO_2_ conditions. (A) Principal Component Analysis (PCA) plot visualizing expression patterns of VLC-inducible genes in each sample. The plot shows the first two principal components (PC1 and PC2), respectively. Samples are color-coded and shape-coded by strain (WT, *ccm1*-1, *cbp1*-1) and CO_2_ condition (HC, VLC 0.3 h, VLC 2.0 h). PC1 accounts for 62.6% of the variance and PC2 for 16.4%. (B) Venn diagram representing the number of differentially expressed genes (DEGs) in *cbp1*-1 under different CO_2_ conditions. Numbers in black represent DEGs specific to or shared between HC, VLC 0.3 h, and VLC 2.0 h conditions. The number of VLC-inducible and CCM1-dependent genes is shown in red numbers. (C) Volcano plots showing Differentially Expressed Genes (DEGs) in *cbp1*-1 mutants acclimated to HC conditions. Left panel: *cbp1*-1 vs WT; Right panel: *cbp1*-1 vs *cbp1*-1:*CBP1*. The x-axis shows log_2_ fold change and the y-axis shows –log_10_(FDR). Red dots indicate Ci transporters, carbonic anhydrases and the transcription factor involved in CCM which are highlighted and labeled. Dashed lines indicate the threshold for significance (FDR < 0.01, |log_2_FC| > 1). (D) Heatmap showing the expression patterns of 27 VLC-inducible and CCM1-dependent DEGs that are mis-regulated in HC condition across different strains (WT, *ccm1*-1, *cbp1*-1) and CO_2_ conditions (HC, VLC 0.3 h, VLC 2.0 h). Gene expression levels are represented as Z-scores of log_10_(TPM) values. Gene IDs and their corresponding gene names are shown on the right. Hierarchical clustering was performed on both genes (rows) and conditions (columns).

Next, we classified differentially expressed genes (DEGs) as CCM1-dependent or CBP1-dependent if their expression in the mutant (either *ccm1*-1 or *cbp1*-1) differed significantly from both WT and the respective complemented strain under any condition (FDR < 0.01, |log_2_ fold change| > 1). Of the 368 CBP1-dependent DEGs identified, 163 were affected under HC conditions, while 244 and 116 showed altered expression at VLC 0.3 h and 2.0 h, respectively (Fig. 4B). The 41 CCM1-dependent DEGs showed the expected pattern of VLC-specific induction. By contrast, CBP1-dependent DEGs were distributed across all conditions and showed no strong clustering at a specific time point (Fig. 4B).

Notably, among the VLC-inducible, CCM1-dependent genes (41 total), 27 were aberrantly upregulated in *cbp1*-1 under HC (Fig. 4B), indicating that CBP1 normally represses their induction when CO_2_ is abundant. Volcano plot analysis confirmed substantial derepression of key CCM genes under HC conditions (Fig. 4C). These include carbonic anhydrases (CAH1, LCIB), HCO_3_^−^ channels (LCIA, LCI1, BST1, BST3), and the transcription factor LCR1 (Fig. 4C and Table S6). The significant upregulation of *CAH1* is likely a downstream consequence of the strong induction of *LCR1*, a Myb transcription factor known to directly activate *CAH1* expression (7). Complementation analysis (*cbp1*-1 vs *cbp1*-1:*CBP1*) confirmed that reintroduction of *CBP1* restored repression of these CCM genes under HC conditions, validating that the observed derepression is specifically due to CBP1 loss. It is also of note that *CAH1* showed enhanced suppression in the complemented strain (Fig. 4C, right panel).

Hierarchical clustering of the 27 mis-regulated genes revealed coordinated upregulation in *cbp1*-1 under HC, with Z-scores ranging from 0 to 2, indicating substantial derepression (Fig. 4D). We also observed upregulation of light stress-related genes (*LHCSR3.1* and *LHCSR3.2*) and a mitochondria-localized membrane protein (*CCP2*) in *cbp1*-1 under HC. The heatmap demonstrates that these genes maintained normal VLC-induced expression in *cbp1*-1, suggesting CBP1 specifically functions in HC repression rather than VLC activation (Fig. 4D and Table S6). Under VLC, however, the expression of these genes in *cbp1*-1 closely resembled WT. We also did detect altered expression of several unknown genes in *cbp1*-1 under VLC (Table S7), suggesting additional functions for CBP1. Collectively, these results indicate that CBP1 acts primarily as a repressor of CCM-related genes under HC conditions.

### Inorganic carbon affinity is increased in *cbp1* mutants acclimated to HC conditions

To investigate whether the transcriptional derepression observed in *cbp1*-1 leads to functional consequences, we examined the accumulation of key CCM proteins. Immunoblot analysis revealed that under HC conditions, *cbp1*-1 showed increased levels of CAH1 and LCIB compared to WT, consistent with our RNA-seq data (Fig. 5A). This elevated protein accumulation was reversed in the complemented strain (*cbp1*-1:*CBP1*), confirming that CBP1 directly regulates these CCM components. Other CCM proteins, including LCIA, LCI1, LCIC, and HLA3, showed similar levels across all strains under HC conditions. Under VLC, all strains exhibited robust induction of CCM proteins, with *cbp1*-1 showing slightly reduced levels of HLA3 and LCI1, though this was not restored by complementation and thus appears CBP1-independent.

**Fig. 5.**
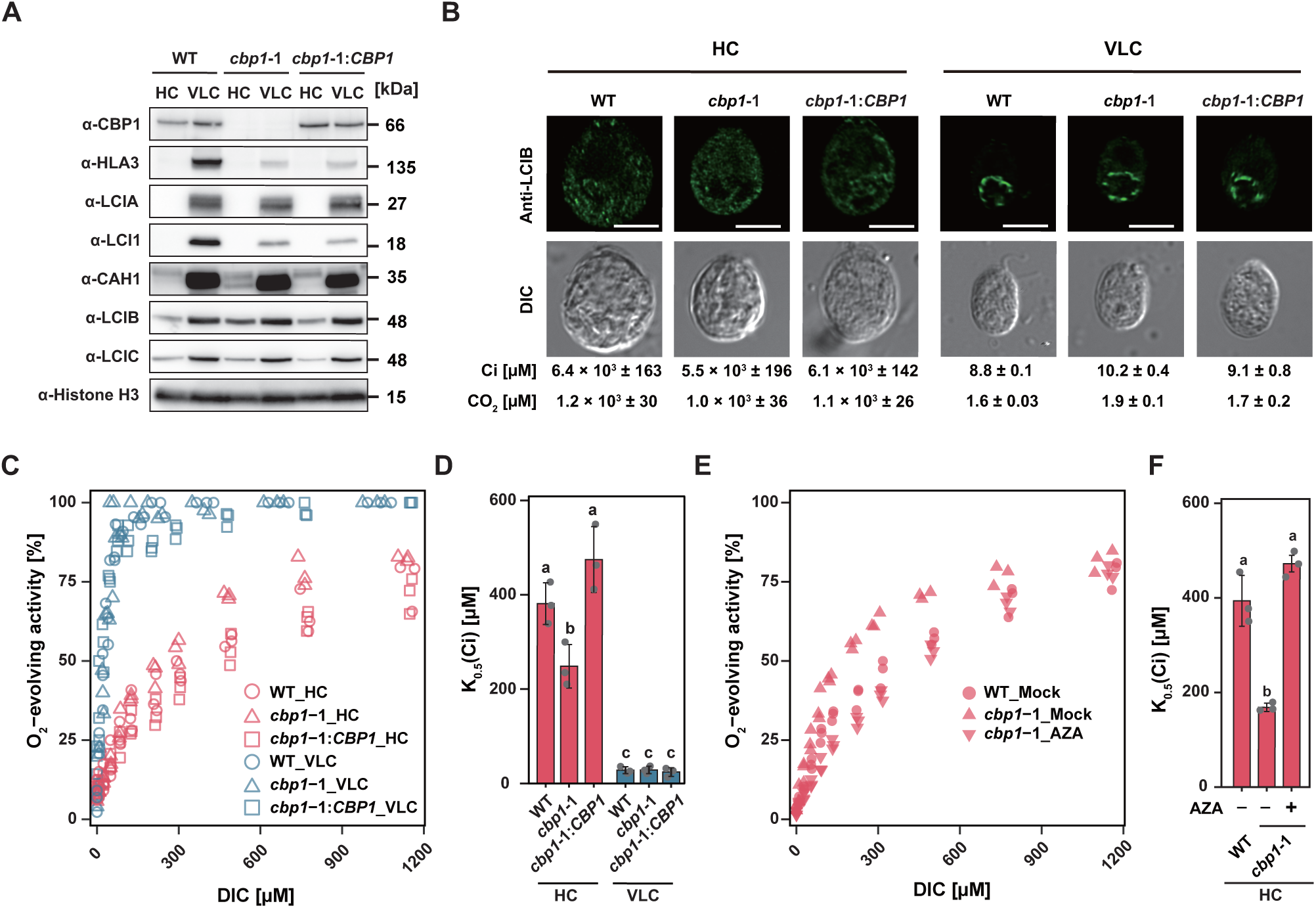
CBP1 loss enhances apparent CO_2_ affinity through elevated carbonic anhydrase activity. (A) Accumulation of CCM-related proteins in WT, *cbp1*-1 and *cbp1*-1:*CBP1*. Cells were first grown under 5% (v/v) CO_2_ condition for 24 h and shifted to 5% (v/v) CO_2_ (HC) or 0.04% (v/v) CO_2_ (VLC) conditions for 12 h. Western blot analysis was performed using antibodies against the indicated proteins. Histone H3 was used as a loading control. (B) Subcellular localization of LCIB in WT, *cbp1*-1, and *cbp1*-1:*CBP1* strains acclimated to HC and VLC conditions. Anti-LCIB immunofluorescence (green) and DIC images are shown. The concentration of inorganic carbon (Ci) in the culture medium and the calculated CO_2_ concentration are shown. Scale bars represent 5 μm. (C) Net O_2_-evolving activities of WT, *cbp1*-1, and *cbp1*-1:*CBP1* cells grown in HC or VLC conditions for 12 h measured against dissolved inorganic carbon (DIC) concentrations at pH 7.8. Data points represent mean values from three biological replicates. (D) K_0.5_(Ci) values representing the DIC concentration required for half-maximal O_2_ evolution in WT, *cbp1*-1, and *cbp1*-1:*CBP1* cells grown in HC or VLC conditions for 12 h. Data represent mean values ± standard error (SE) from three biological replicates. Statistical analysis was conducted using the Tukey–Kramer multiple comparison test, with different letters indicating significant differences (P < 0.05). (E) Net O_2_-evolving activities of WT and *cbp1*-1 cells treated with or without acetazolamide (AZA) grown in HC conditions for 12 h measured against DIC at pH 7.8. Mock treatment with 1% DMSO was used as vehicle control. (F) K_0.5_(Ci) values of WT and *cbp1*-1 treated with or without AZA. AZA was adjusted to a concentration of 5 mM, dissolved in DMSO, and added to the measuring buffer at 1% (v/v). Data represent mean values ± SE from three biological replicates. Statistical analysis was performed using the Tukey–Kramer multiple comparison test, with different letters indicating statistically significant differences (P < 0.05).

**Fig. 6.**
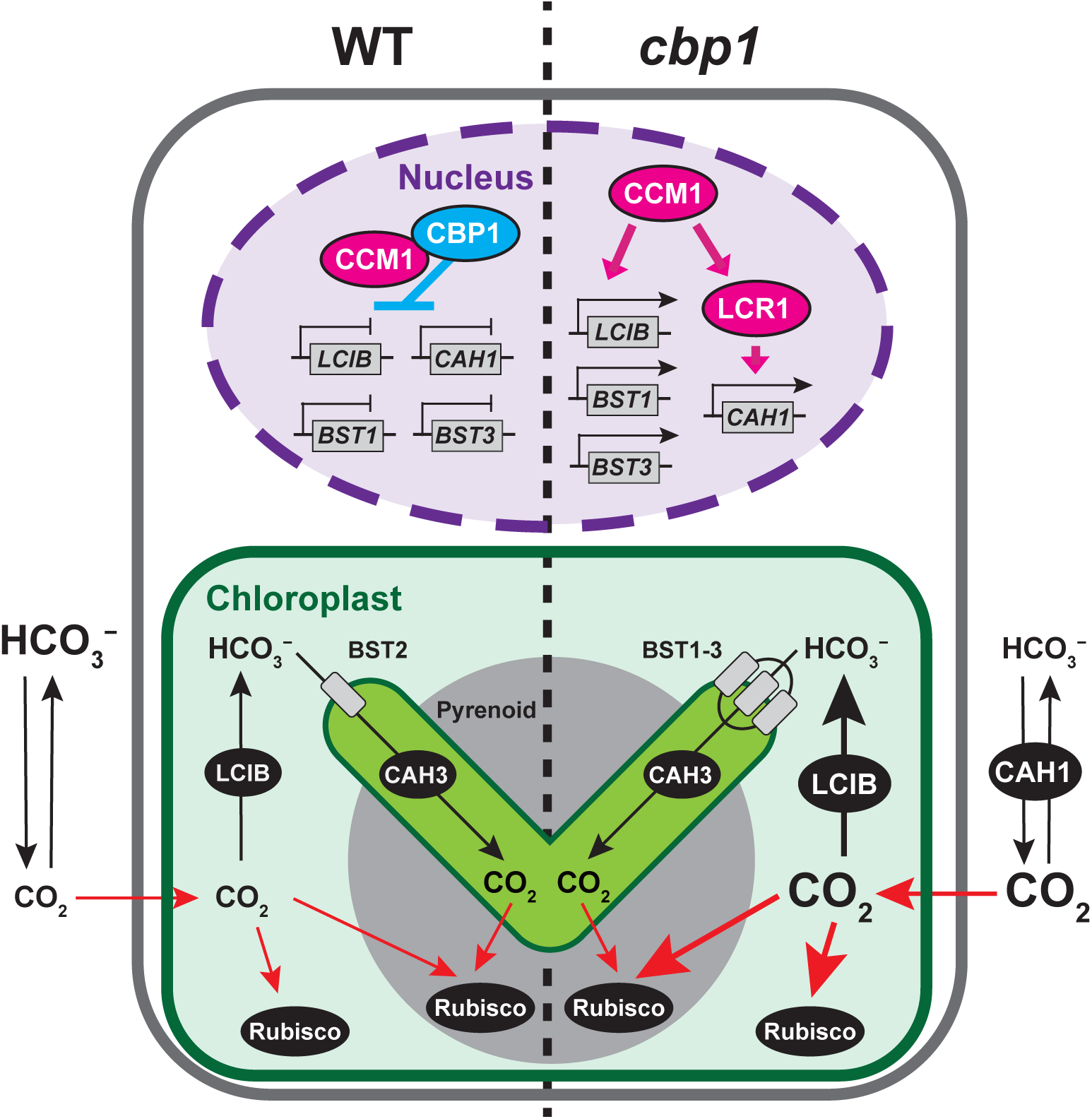
Model illustrating the regulatory role of CBP1 in suppressing CCM gene expression under high CO_2_ conditions. Under HC conditions, the model compares CCM regulation between wild-type (WT, left) and *cbp1* mutant (right) cells. In WT cells, CBP1 physically interacts with CCM1 in the nucleus, preventing CCM1 from activating transcription of CCM genes including *BST1*, *BST3*, *LCIB*, and *CAH1* (indicated by repression bars on gene promoters). This results in minimal expression of CCM components, with only basal levels of carbonic anhydrases (LCIB in chloroplast stroma and CAH3 in thylakoid membranes). CO_2_ passively diffuses across membranes (red arrows) to reach Rubisco. In *cbp1* mutant cells (right), the absence of CBP1-mediated repression allows CCM1 to initiate a complex transcriptional cascade even under HC conditions. This cascade includes the activation of the transcription factor LCR1, which in turn induces its direct target, *CAH1*. In parallel, other key CCM genes such as *BST1*, *BST3*, and *LCIB* are also activated (shown by active transcription from gene promoters), although whether this is due to direct regulation by CCM1 or mediated by other unidentified transcription factors remains unknown. This widespread derepression leads to the increased accumulation of CAH1 and LCIB proteins, as confirmed by immunoblot analysis (Fig. 5A). The elevated CAH1 in the periplasmic space enhances the dehydration of HCO_3_^−^ to CO_2_. This increased carbonic anhydrase activity enhances the local CO_2_ concentration at the cell surface, improving CO_2_ capture efficiency and thereby explaining the enhanced apparent Ci affinity observed in *cbp1* mutants. The BST1-3 proteins, which function as HCO_3_^−^ transporters, also show increased expression.

We next examined LCIB localization, as its redistribution from diffuse to punctate pattern around pyrenoid is a hallmark of CCM activation. Despite the increased LCIB protein levels in *cbp1*-1 under HC, its localization remained diffuse in all strains under HC conditions (Ci: 5.5–6.4×10^3^ µM, CO_2_: 1.0–1.2×10^3^ µM), similar to WT (Fig. 5B). Under VLC conditions (Ci: 8.8–10.2 µM, CO_2_: 1.6–1.9 µM), LCIB formed distinct puncta in all strains, indicating that CBP1 loss does not affect LCIB relocalization dynamics.

To assess whether increased CAH1 and LCIB levels enhance CO_2_ utilization, we measured photosynthetic O_2_ evolution across a range of dissolved inorganic carbon (DIC) concentrations. The *cbp1*-1 mutant acclimated to HC showed a leftward shift in its DIC response curve compared to WT, indicating enhanced apparent Ci affinity (Fig. 5C).

Quantification revealed that K_0.5_(Ci), the DIC concentration supporting half of maximal O_2_ evolution (V_max_), decreased from 381 ± 26 µM in WT to 248 ± 27 µM in *cbp1*-1 under HC conditions (P < 0.05), while complementation restored it to 475 ± 40 µM (Fig. 5D and Table S5). Under VLC conditions, all strains showed similarly low K_0.5_(Ci) values (∼20-30 µM), confirming that the enhanced affinity phenotype is HC-specific. V_max_ were comparable across all strains under both conditions (Table S5).

To determine whether the enhanced Ci affinity in *cbp1*-1 results from elevated CA activity, we treated cells with acetazolamide (AZA), a membrane-impermeable inhibitor of extracellular CAs including CAH1. AZA treatment (5 mM) increased K_0.5_(Ci) in HC-acclimated *cbp1*-1 from 169 ± 5 µM to 416 ± 70 µM, effectively restoring it to WT levels (394 ± 31 µM) (Fig. 5E and F and Table S5). This complete rescue by CA inhibition demonstrates that excess CAH1 activity is the primary driver of enhanced Ci affinity in *cbp1*-1.

Collectively, these results reveal that CBP1 maintains appropriate CCM suppression under HC conditions by preventing premature accumulation of CCM proteins, particularly CAH1. Loss of this regulatory control leads to constitutive CA activity that enhances CO₂ capture efficiency even under carbon-replete conditions, potentially representing an energetic burden to the cell.

## Discussion

In this study, we identified the long-postulated nuclear “off switch” that deactivates the algal CCM when CO_2_ becomes abundant. By screening for CCM1 interactors, we discovered CBP1, a CobW/WW-domain protein that localizes to the nucleus, binds CCM1 constitutively, and is required to repress a defined subset of CCM1-dependent genes under HC conditions. This finding addresses a significant gap in our understanding of how green algae deactivate the CCM and establishes a Zn-linked regulatory layer that complements the LysR-type and bZIP-type repressors known in cyanobacteria and diatoms.

### Zn-Linked Control of CCM1 Activity

CBP1 belongs to the GTP-driven COG0523 family and thus almost certainly transports divalent metals (20–22). This is highly significant because CCM1 itself requires two tightly bound Zn^2+^ ions for structural integrity and transcriptional activity: mutating any of the coordinating residues C36, C41, H54, C77 or C80 eliminates Zn binding, destabilizes the 290–580 kDa CCM1 complex and prevents CCM induction (13, 14). Based on these findings, we propose a working hypothesis, the ‘Zn-switch model’, in which the CobW/CobW_C domain of CBP1 may modulate the Zn^2+^ occupancy of CCM1, potentially toggling the regulator between active and inactive states, while the WW domain— functionally similar to YAP/TAZ or BES1 scaffolds (26–28)—could recruit auxiliary repressors. This model provides a compelling framework for understanding how CCM1 activity could be rapidly toggled. However, demonstrating direct zinc transfer between CBP1 and CCM1 and its functional consequences on transcriptional activity are essential future objectives required to validate this hypothesis.

BiFC confirms CBP1-CCM1 binding, yet only 27 of 41 CCM1 targets are de-repressed in the *cbp1* mutant, suggesting additional nuclear clients yet to be identified. Furthermore, the relatively limited impact of CBP1 disruption under HC conditions strongly indicates the existence of additional repressors, potentially among the other two CobW/WW-domain proteins closely related to CBP1 identified in our phylogenetic analysis (Fig. 1D). Functional redundancy or cooperative interactions among these proteins could collectively ensure robust and precise CCM repression in response to elevated CO_2_. Future studies dissecting the roles of these paralogs will further clarify the complexity and modularity of CCM repression mechanisms. Demonstrating Zn transfer and mapping CBP1 chromatin occupancy by ChIP-seq are also essential next steps.

### CBP1-Dependent Genes and Potential Regulatory Mechanisms

Loss of CBP1 triggers a complex transcriptional cascade under HC conditions, revealing new layers of CCM regulation. Part of this cascade follows a known pathway: the derepression of the Myb transcription factor LCR1 leads to the subsequent activation of its direct target, *CAH1* (7). However, the transcriptional activators for other key derepressed genes, such as LCIB and the BST1/3, are not downstream of LCR1 and have remained elusive. Our data show that CBP1 also represses several other previously unrecognized genes, suggesting their involvement as novel CCM components or modulators. For instance, LCI9 and the methyltransferase SMM7 may have roles in pyrenoid assembly or maintenance (29). More importantly, two additional transcription factor genes, *Cre03.g212977* and *Cre03.g212641*, are also derepressed in the *cbp1*-1 mutant (Table S6). These factors, which respond to both low CO_2_ and cold stress (30), are strong candidates for the missing activators that control *LCIB, BSTs*, and other CCM genes in parallel to the CAH1-LCR1-CAH1 axis, potentially via chromatin remodeling (31). The discovery of this complex network, governed by a protein with a GTP-dependent zinc transferase domain like CBP1, is intriguing. Functional precedents in yeast and vertebrates link Zn trafficking directly to transcriptional regulation (32, 33), raising the possibility that CBP1 modulates this network via zinc-dependent control of transcription factor activities or chromatin structure. Therefore, further characterization of these CBP1-regulated pathways will be required to understand how zinc metabolism, transcriptional regulation, and stress adaptation are integrated to finely regulate CCM activity.

### Alternative metal-swap scenarios for CCM suppression

COG0523 proteins can mismetallate their clients when intracellular metal ratios shift. *E. coli* CobW, for example, binds Zn^2+^ more tightly than Co^2+^ in vitro but delivers Co^2+^ *in vivo* only when the cytosolic Co/Zn ratio is high (34). Analogously, CBP1 could insert Mn^2+^ or Co^2+^ into the Zn sites of CCM1 under HC conditions, rendering CCM1 inactive until Zn^2+^ again predominates. A second, non-exclusive possibility is simple Zn scavenging: if CBP1-GTP binds Zn^2+^ with sub-nanomolar affinity, it could extract the metal from CCM1, leaving an apo form that rapidly loses activity. These hypotheses are testable by (i) quantifying metal-binding hierarchies of CBP1-GTP versus CCM1, (ii) measuring real-time ^65^Zn transfer between the two proteins in the presence of competing Mn^2+^/Co^2+^, and (iii) assaying the transcriptional competence of mismetallated CCM1 in heterologous reporter systems. Whether CBP1 acts as a co-activator, co-repressor or Zn chaperone will ultimately depend on such biochemical evidence.

### A putative link between carbon-nitrogen status and CCM control

Besides CBP1, our pull-down consistently retrieved glutamate dehydrogenase GDH1/2 (Fig. 1A and Table S1–4), an enzyme that reversibly converts glutamate and α-ketoglutarate while shuttling NAD(P)H (35). GDH1/2 was lost after 1.5 M KCl washes (Fig. 1A) and, unlike CBP1, we failed to isolate GDH1/2 mutants despite four independent CRISPR guides and screening > 1,000 colonies, echoing earlier biochemical work showing that all three NAD(P)^+^-dependent GDH isozymes are mitochondrial and indispensable for amino-acid catabolism in *Chlamydomonas* (36). We speculate that GDH1/2 docks only transiently on CCM1 to couple cellular C/N or redox status to CCM shutdown: when external CO_2_ is high, a rising glutamate:α-ketoglutarate ratio would favor nitrogen assimilation and could provide an additional signal to disengage the CCM. Determining whether GDH1/2 alters CCM1 post-translationally, or whether its metabolic products modulate the CBP1 Zn-switch, will require conditional GDH knock-down and targeted metabolomics, but the weak yet reproducible interaction uncovered here points to a promising carbon-nitrogen crosstalk layer for future study.

### Physiological impact, evolutionary footprint and applied outlook

Although a single spot-test plate (Fig. 2D) suggested slightly reduced colony expansion of the cbp1-1 mutant at 5 % CO_2_, this effect was not reproducible in independent growth assays in liquid or on solid medium; thus partial CCM derepression under carbon-replete conditions does not confer a measurable growth disadvantage. We speculate on several reasons for this mild phenotype. First, the energetic burden of this partial derepression, while real, may be too small to impact fitness under nutrient- and light-replete laboratory conditions; it is plausible that a growth phenotype would emerge under more ecologically relevant, competitive scenarios where energy resources are limited. Second, the observed derepression of CCM genes in *cbp1*-1 is still considerably lower than their full induction levels under VLC conditions, suggesting the actual energetic cost may be modest. Furthermore, the existence of potential functional redundancy among the CBP1 paralogs identified in our phylogenetic analysis could also contribute to the mild phenotype, as these related proteins might partially compensate for the loss of CBP1.

In contrast, the *ccm1*-1 mutant, which is completely unable to induce the CCM, consistently exhibits slightly better growth than wild-type under HC conditions (Fig. 2D). This suggests that wild-type cells incur a fitness cost from maintaining even a basal level of the CCM machinery. Indeed, previous transcriptome analyses have established that many CCM component genes maintain a low but distinct basal level of expression in wild-type cells even under HC conditions (11). The complete absence of this basal expression in *ccm1*-1 likely provides a net energy saving that promotes growth when CO_2_ is not limiting. Taken together, the contrasting phenotypes of *cbp1*-1 and *ccm1*-1 highlight the delicate energetic trade-offs associated with CCM regulation and the importance of its strict suppression when CO_2_ is abundant.

The co-occurrence of WW-domain COG0523 proteins and CCM1 exclusively in Chlorophyceae and Trebouxiophyceae suggests that the CCM1-CBP1 module is a lineage-specific solution for reversible CCM control, distinct from the LysR- and bZIP-type repressors used by cyanobacteria and diatoms (15–17). Whether higher plants employ a comparable Zn-based switch for their CO_2_ sensors remains an open and exciting question (2). From an applied perspective, modulating CBP1 activity offers a strategy for metabolic engineering: attenuating CBP1 could enhance CO_2_ capture from flue gas, while enhancing it might conserve ATP in carbon-rich photobioreactors, provided that zinc homeostasis is maintained. By positioning CBP1 at the nexus of metal metabolism and transcriptional repression, this study offers both a mechanistic framework for nuclear metal signaling and a blueprint for engineering algal CO₂ fixation.

## Materials and methods

### *Chlamydomonas reinhardtii* strains and culture conditions

We used two *Chlamydomonas reinhardtii* strains in this study: (1) a wild-type strain designated C9 (originally from the IAM Culture Collection, University of Tokyo; currently maintained as NIES-2235 and CC-5098), and (2) the *ccm1* mutant (25). Unless otherwise noted, cells were grown in Tris-acetate-phosphate (TAP) medium at 25°C under continuous illumination (∼120 μmol photons m⁻² s⁻¹) with gentle shaking (120 rpm). Once cultures reached mid-logarithmic phase (optical density at 730 nm [OD₇₃₀] = 0.4–0.7), cells were harvested by centrifugation (2,000 g, 5 min) and transferred to MOPS-buffered phosphate (MOPS-P) medium for experimental treatments.

For high-CO_2_ (HC) conditions, cultures were bubbled continuously with 5% (v/v) CO_2_ in air, whereas for very low-CO_2_ (VLC) conditions, cultures were aerated with ambient air (∼0.04% CO_2_). Both conditions were maintained at 25°C with the same light intensity (∼120 μmol photons m⁻² s⁻¹). When necessary, we monitored growth by measuring OD_730_. All experiments were performed on cells in the mid-logarithmic growth phase to ensure consistent physiological states.

### Pull-down assay

Cells were grown photoautotrophically in MOPS-P medium under the appropriate CO_2_ conditions (HC or VLC) until OD_730_ = 0.4–0.7. Typically, 1–3 L of culture was harvested at 4°C by centrifugation (3,000 g, 5 min). Pellets were resuspended in suspension buffer (300 mM Tris-HCl [pH 7.5], 300 mM KCl, 15 mM MgCl_2_) supplemented with 3×protease inhibitor cocktail (EDTA-free, Roche). Cells were disrupted by sonication (Handy Sonicator UR-20P, TOMY; amplitude setting 8, 8 cycles of 5 s ON/3 s OFF on ice).

Lysates were clarified by sequential centrifugation at 10,000 g for 10 min and 100,000 g for 20 min at 4°C, followed by filtration through a 0.22 µm membrane. The protein concentration was determined by Bradford assay and adjusted to 5 mg mL⁻¹. For the pull-down, M2 FLAG Affinity Gel (Sigma-Aldrich) was pre-equilibrated in base buffer (100 mM Tris-HCl [pH 7.5], 5 mM MgCl_2_, 0.1% Triton X-100). The lysate (typically 10 mg total protein) was incubated with the gel (4 µL slurry per 10 mg protein) at 4°C for 2 h with gentle rotation. After binding, the gel was washed five times each with buffers of increasing ionic strength (100, 500, and 1,500 mM KCl in base buffer). Bound proteins were eluted with 500 ng µL⁻¹ of 3×FLAG peptide (Sigma-Aldrich) in base buffer by overnight rotation at 4°C. Eluates were collected by centrifugation (500 g, 1 min, 4°C) and pooled for downstream analyses (SDS-PAGE and mass spectrometry).

### Mass spectrometry analysis

Protein bands were excised under blue light using a Safe Imager Blue-Light transilluminator (Invitrogen). Gels previously visualized by FLA3000 were placed on the illuminator, and regions of interest were excised with a surgical blade. Mass spectra were obtained from the gel pieces excised and subsequently digested with trypsin using an LC-MS/MS spectrometer (LTQ: Thermo Fischer Sciencific) (37). Protein identification was performed by peptide mass fingerprinting using BioWorks software (Thermo Science).

### Construction of BiFC vectors

To visualize protein-protein interactions by bimolecular fluorescence complementation (BiFC), cDNA fragments corresponding to *CCM1* and *CBP1* were cloned into Gateway entry vectors (Thermo Fisher Scientific) using In-Fusion HD Cloning (Takara). Each gene was fused in-frame with either the N-terminal (nEYFP) or C-terminal (cEYFP) fragment of YFP. LR recombination was subsequently performed to generate destination vectors for transient expression.

### Agrobacterium-mediated *Nicotiana benthamiana* leaf infiltration

*Agrobacterium tumefaciens* strain carrying the BiFC constructs was grown in YEP medium containing the appropriate antibiotics. Bacterial cells were harvested by centrifugation, resuspended in infiltration buffer to OD_600_ = 0.8, and incubated at room temperature for 1–2 h. The suspensions were syringe-infiltrated into the abaxial side of *N. benthamiana* leaves according to Miyaji et al. 2025 (38). Two to three days post-infiltration, small leaf sections were excised and observed under a confocal laser scanning microscope (see below).

### Subcellular localization analysis of CCM1 and CBP1

To observe the subcellular localization of CCM1 and CBP1 in *Chlamydomonas*, genomic regions of CCM1 and CBP1, including their ∼2.0 kb upstream sequences, were first cloned into pENTR™ 1A Dual Selection Vector (Thermo Fisher Scientific) using In-Fusion^®^ HD Cloning Kit (Takara Bio). These entry clones were then recombined with destination vectors containing mGold through Gateway LR reactions (Thermo Fisher Scientific) to generate expression constructs. The resulting constructs were transformed into the relevant mutant backgrounds (e.g., *ccm1*-1 or *cbp1*-1) by electroporation (39).

Transgenic cells were grown under either HC or VLC conditions, then harvested at mid-log phase. For fluorescence microscopy, cells were placed on 1.0% agarose pads and sealed under coverslips. Confocal images were acquired using a Leica SP8 confocal microscope with a 63×oil-immersion objective. mGold fluorescence was excited at 543 nm, and emission was detected between 560 and 600 nm. Images were processed in LAS X software (Leica) for minor adjustments of brightness and contrast.

### Measurement of photosynthetic O**_2_**-evolving activity

Cells were harvested by centrifugation (1,000 g, 5 min, 25°C) and resuspended in Ci-depleted 20 mM HEPES–NaOH (pH 7.8) at a final chlorophyll concentration of 10–20 µg mL⁻¹. Oxygen evolution rates were measured at 25°C using a Clark-type oxygen electrode (Hansatech Instruments) under saturating white light (∼700 µmol photons m⁻² s⁻¹) as described previously (40). NaHCO_3_ was injected stepwise to vary the Ci concentration. K_0.5_(Ci) values, defined as the Ci concentration required for half-maximal O₂ evolution (V_max_), was calculated by fitting the data to the Michaelis–Menten equation.

For acetazolamide (AZA) treatments, AZA was dissolved in DMSO at 5 mM and added to the cell suspension at a final concentration of 1% (v/v). Control samples received 1% DMSO only.

### CRISPR-Cas9 system-mediated generation of mutants

To generate *cbp1*-1, we designed a guide RNA (5′-TGAGTTCTGAACGTTTGTCGGGG-3′) targeting the second exon of the *CBP1* gene using CRISPOR (http://crispor.tefor.net/) (41). The crRNA and tracrRNA were chemically synthesized (Integrated DNA Technologies) and mixed with recombinant Cas9 protein to form a ribonucleoprotein (RNP) complex. RNP and the *AphVII* cassette (conferring hygromycin resistance) were co-introduced into WT C9 cells by electroporation (42).

Transformants were selected on TAP plates containing 30 µg mL⁻¹ hygromycin and screened by colony PCR. Specific genomic insertion of the *AphVII* cassette was verified by PCR with primers 5′-GTTGTAGGTGGGTTGGAGGG-3′ (forward) and 5′-ATGTCCTTCGCCTTCTCAGC-3′ (reverse). Immunoblotting with a CBP1-specific antibody confirmed the loss of CBP1 protein in the resulting mutant.

### Immunoblot analysis

Harvested cells (OD_730_ = ∼0.6) were pelleted by centrifugation at 2,000 g for 5 min, then resuspended in SDS loading buffer (50 mM Tris-HCl [pH 8.0], 25% glycerol, 2% SDS, 100 mM DTT). After incubation at 37°C for 30 min, lysates were clarified by centrifugation (13,000 g, 10 min). Approximately 10 µg of total protein per lane was separated on SDS-PAGE and transferred onto a PVDF membrane (Fluoro Trans, Pall Life Science) using a semidry blotter.

Membranes were blocked in 5% (w/v) skim milk (Wako) in PBS at room temperature for 1 h, followed by washing in PBS containing 0.1% Tween-20 (PBS-T). The following primary antibodies were used: anti-HLA3 (1:1,250), anti-LCIA (1:5,000), anti-LCI1 (1:5,000), anti-LCIB (1:5,000), anti-CBP1 (1:5,000), anti-CAH1 (1:2,500), anti-CAH3 (1:2,000), anti-CCM1 (1:2,500), and anti-Histone H3 (1:20,000). HRP-conjugated goat anti-rabbit IgG (Life Technologies) was used as secondary antibody at 1:10,000 dilution.

### RNA-seq analysis

Total RNA was isolated using the RNeasy Plant Mini Kit (QIAGEN) and sequenced on an Illumina NovaSeq 6000 platform with biological duplicates for each condition. Sequence reads were mapped to the *Chlamydomonas reinhardtii* genome (v5.6, Phytozome). The alignment, read counting, and Trimmed Mean of M-values (TMM) normalization of read counts were performed according to the methods previously described in Shimamura et al. 2023.

## Data Availability

Data deposition: The raw RNA-seq data generated in this study have been deposited in the DDBJ Sequence Read Archive under BioProject accession PRJDB17792, with Run accession numbers DRR635765–DRR635812.

## Accession numbers

The accession numbers of the Phytozome database for *Chlamydomonas* genes *CBP1* is *Cre16.g684650*.

## Acknowledgments

We thank Yu Sasaki and Kentaro Azuma for their technical assistance in conducting the experiments, and Dr. Taiho Kambe for helpful discussion.

