## Supplemental figures and tables for "A nuclear CobW/WW-domain factor represses the CO_2_-concentrating mechanism in the green alga *Chlamydomonas reinhardtii*"

### Fig.S1

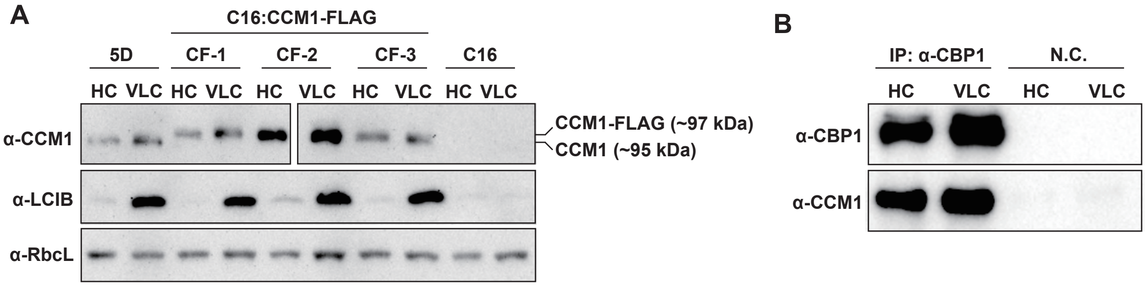

**Fig.S1. Generation of CCM1-FLAG lines for MS analysis and verification of the CCM1–CBP1 interaction.**
**(A)** To select a functionally complemented line for subsequent analysis, the CCM1-FLAG construct was introduced into the ccm1 mutant strain (C16). Three independent transformants, designated CF-1, CF-2, and CF-3, were analyzed by immunoblotting. Functional complementation was confirmed by the restored accumulation of the CCM component LCIB under very-low-$CO_2$ (VLC) conditions. Among these lines, CF-2 showed the highest accumulation of both CCM1-FLAG and LCIB and was therefore selected for the pull-down assay. Cells pre-grown at 5 % $CO_2$ were shifted for 12 h either to the same high-$CO_2$ condition (HC, 5 %) or to very-low-$CO_2$ air (VLC, 0.04 %). RbcL served as the loading control. **(B)** Co-immunoprecipitation of CCM1 with CBP1. Total protein extracts from cells grown as in (A) were subjected to immunoprecipitation (IP) with anti-CBP1 antibodies; precipitates were analysed by immunoblotting with anti-CCM1 and anti-CBP1. "N.C." denotes the negative control in which no antibody was added to the IP reaction.

#### Fig.S2

**>CCM1-FLAG (721 aa)**

1 **MEALDAQDSL QLDVVSPSAR PAAAGGDKR**D PERFYCPYPG CNR**SFAELWR** LKVHYRAPPD
61 IRGSGKERGH GTELTHCPKC GKTLKPGKHH VGCSGGKSAP RQTASKRNR**T GADDADEAVP**
121 **GSPHSK**HVRG TDMDGDPHK**S WQDFALTHAG YAIGAPAMLA PLKQEHPEWP PTVPQGVFVG**
181 **HGDRVSWLPG QVNGFVPQLQ PQR**YQQPQFP PELAQAFAAA GTHAPHVYAQ QVPFASIPGY
241 PGQPGVATLQ VTTESGQVLS IPANMAGMPP GMAGLPGTLV YHQQPPPHDA AASYLAQAQA
301 HAQHAAAMHA VNSAHAQQQQ QQQQQQQQQQ PGVPAAPPAV PGVHDGMPPG TVAAAAAAAA
361 AAAAVGGSAP SALQTDVGGR **PGAALPPQAA PGTGAGQGAG APAGAADGGA APAAGDAAAS**
421 **GGAKPVADED NLGTVFDDVE EFTRDFGRIP SPPPLPPDFH TAATGGNGML FNFSQFGQK**L
481 PRTQSHTRLD RSLSAVGLGH LDVGVDGDVM YDHTDDGDLM QLLFGVPDEL PTMATIHLHK
541 WSNEEDEDDD AAEPGGGGAA AAGGGGGAAA GAGGEGGGGA GAGGGGAGAG AGEANAAAGR
601 GGAGPGPGLE AGGGGGGGGA GEGGPGAGQQ PPHHQQSVGG HDQRPLNGKT **LHGHDASLAV**
661 **LPAPGGKSLM NGGAGHAGEE HHRDHLLDAE TFR**LLQSCDD YKDHDGDYKD HDIDYKDDDD
K

**>CBP1 (606 aa)**

1 MASAGATAPT NVQNSQK**TPV TIITGFLGAG K**TTLLNYILK EKGSR**SIAVI ENEFGEVNID**
61 **RELVAANLLA K**EDLVSLENG CVCCSLRKDI VKAFAEIERR SRQAGGK**KVD AIVLETTGLA**
121 **DPAPVAFTFF ANPWIASR**FR **LDSIICVVDA RYLMQHLEDG KHSDGTVNEA VQQIAFADLI**
181 **LLNKIDLVNS EEQKK**QVLGA IRAVNNSARI **VECQLNQETG RPHMDMLLFN NLFSVNR**VLE
241 QIDPQFLDSD SDDDAEDEEE APSPQSGTQA QQDQASRAAK **GQAAEPGAAA GADAGAGPGS**
301 **K**ATAEASAGS KAGAGCCGSG EGKGK**AAVED EAAARTAGEA GPSGSSAADA K**AKLLQK**YAD**
361 **KGSVAGHK**HT RDDNCEDNCE ECHIVDGMPI KGERNPKRRA K**RLHDLSDVS SVGIMAR**GPL
421 DEYRFNMYMR DLLAEKAKDI FRCK**GVLSVH GYGSTK**FVFQ GVHETICYGP AEQPWKPEEQ
481 R**VNQVVFIGR** GLNR**KALIEG FR**TCVWVPLP DGWDEFRDTT TKQPFYVNRN TGEKSWTRPE
541 IACARVVATQ GK**TQQPSQLL PRRTASTVGQ LALQQAAAAA ASGAVAPATA SAK**AGGAASA
601 ATEVAS

**>GDH1 (448 aa)**

1 MEARTVSVLA GLAHRVASAA WRSASGLRTC TTTTPTYGRT SVYVK**EALDL LDFDPQVEKA**
61 **ILNPDREVTV NLVVPMDNGE VNMFPAYR**VQ HNNALGPFK**G GIIYHPGVTL ENMR**NLASLN
121 TWK**FSLLNVQ FGGAK**GGVGV DPRSLSERET EKLTRKYVQA LQEVIGPHTD IPAPDINTDE
181 HHMAWIFDQY SRLRGFAPAA VTGKPTWLHG IVGRDKAGGR GAAIATREFL TRSLRRK**VAG**
241 **TSFLIQGFGK LGSWTAQILQ QEMGAK**IVGV SCSETAVYNE EGLDIPALRA HVAAGGLLKD
301 FPGGTGVLND DSFLDLPADV FIPCAVDGTI HAGNVHRCVN FK**AVVEAANG ALTPEADAAL**
361 **RK**AGVPVLPD LIANGGAVVV SFFEWVQNNQ NMQWEEDDVK RELDR**YLTDA FEALLR**EQSL
421 HAGCSLR**TAG YLVALR**RLQQ ADSVRGHS

**>GDH2 (449 aa)**

1 MASALLACGR QLSCALGGLS LGHQALGAIG GAFRRHASSH AENTNTFLRE ALVKLDYPEK
61 **LQNLLLTPRR EMSVELVVQM DDGQIEVFNA YR**VQHNNARG PYKGGLR**YHP QVDLDDVRSL**
121 **ASLMTWKTAV MDIPYGGAK**G GVTVDPRKLS ERELEKMTRK LVVAIK**EIIG TYEDIPAPDM**
181 **NTDAKVMAWF FDEYSK**YK**GF SPGVVTGKPV YLHGSLGR**EA ATGRGTTFAI RELLKALHMG
241 KIADQK**YVIQ GFGNVGAWAA QLLWEAGGK**V VAISDVAGAV HNEQVRGLDI GALR**KHVASG**
301 **KPLAEFTGGA AVPK**QDILLH PCDVLIPAAI GGVIGPEEAK KLQCK**VVVEA ANGPTTPEGD**
361 **MVLR**DR**GITV LPDIYTNGGG VTVSFFEWVQ NLQNFKWEED DVNR**KLDRK**M ADAFAALWAV**
421 **HK**EMNVPLR**T AAFVVALQR**V TRAEVHRGFD

**Fig.S2. Sequence coverage of CCM1, CBP1, GDH1 and GDH2 obtained by LC-MS/MS.**
Full-length amino-acid sequences are shown for each protein, with regions identified by mass spectrometry highlighted in red. Sequence coverage amounts to 37.1 % for CCM1, 53.0 % for CBP1, 31.5 % for GDH1 and 52.3 % for GDH2. In the CCM1 panel, three serine phosphorylation sites are annotated: S122 and S451 (boxed in black) were detected under both high-$CO_2$ and very low-$CO_2$ conditions, whereas S10 (boxed in red) was observed only under very low-$CO_2$ (VLC) conditions. Residues derived from the C-terminal FLAG tag are shown in blue.

### Fig.S3

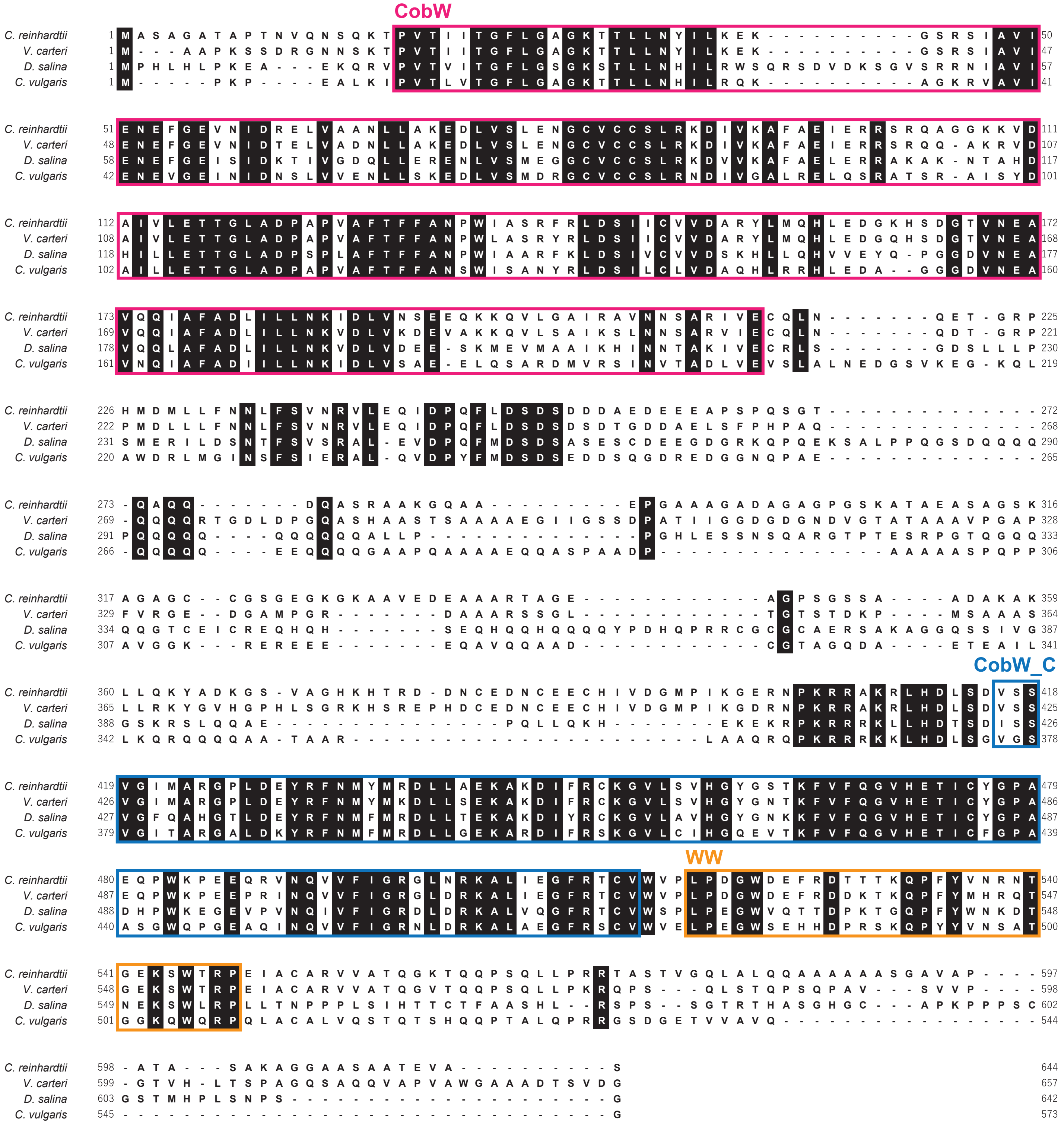

Fig. S3. Multiple sequence alignment of CBP1 proteins from various green algae species. The alignment includes sequences from *Chlamydomonas reinhardtii*, *Volvox carteri*, *Dunaliella salina*, and *Chlorella vulgaris*. Identical amino acids are highlighted in black. The alignment reveals conserved regions across the different species, particularly in the CobW ,CobW_C and WW domains. Numbers on the right indicate the amino acid positions. Gaps in the alignment are represented by dashes. The alignment was performed using MEGAX.

Table S1. Proteins identified by LC-MS/MS from immunoprecipitated gel bands.

| Sample name | | Reference | P (pro) | Sf | Score | Coverage | MW | Peptide (Hits) |
|---|---|---|---|---|---|---|---|---|
| 01_CCM1 | 1 | estExt_fgenesh2_pg.C_50429 {CCM1} regulator of CO2-responsive genes | 7.07E-13 | 7.50 | 80.30 | 21.30 | 70033.89 | 9 (9 0 0 0 0) |
| | 2 | estExt_gwp_1H.C_900015 {ADH1} | 1.02E-08 | 0.94 | 10.17 | | 102168.10 | 1 (1 0 0 0 0) |
| 02_74k | 1 | RWI_chlre3.14.27.2.51 Nickel chaperone for hydrogenase or urease, putative | 3.28E-13 | 14.43 | 150.32 | 34.10 | 64549.51 | 15 (15 0 0 0 0) |
| | 2 | estExt_fgenesh2_pg.C_50429 {CCM1} regulator of CO2-responsive genes | 6.05E-12 | 0.98 | 10.29 | | 70033.89 | 2 (2 0 0 0 0) |
| | 3 | Chlre2_kg.scaffold_1000769 | 5.86E-07 | 0.96 | 10.17 | | 75461.40 | 1 (1 0 0 0 0) |
| | 4 | Chlre2_kg.scaffold_1000770 | 1.66E-07 | 0.95 | 10.16 | | 67702.49 | 1 (1 0 0 0 0) |
| 03_45k | 1 | estExt_GenewiseW_1.C_760022 {GDH2} glutamate dehydrogenase | 4.76E-10 | 3.75 | 40.23 | 9.30 | 49004.55 | 4 (4 0 0 0 0) |
| | 2 | estExt_fgenesh1_pm.C_440001 {IDA5} Actin | 6.30E-06 | 1.89 | 20.19 | | 41808.93 | 2 (2 0 0 0 0) |
| | 3 | fgenesh2_pg.C_scaffold_2000011 | 2.58E-05 | 0.93 | 10.18 | | 44510.41 | 1 (1 0 0 0 0) |

Proteins present in three gel slices—whole CCM1 immunoprecipitate (01_CCM1), the ~75 kDa band (02_74k), and the ~45 kDa band (03_45k)—were excised, digested with trypsin, and analysed by LC-MS/MS. Mass spectra were acquired on a Q-TOF instrument (Waters) at NAIST and on an LTQ ion-trap (Thermo Scientific) at Okayama University, and searched with Mascot or BioWorks against the Chlamydomonas reinhardtii protein database (JGI v3.1). Columns are as follows. Reference: JGI protein ID and annotation returned as the top hit. P (pro): ranking of the hit within the sample. Sf: Mascot/BioWorks scoring function (lower E-value or higher score indicates greater confidence). Score: confidence score assigned by the search programme. Coverage (%): percentage of the predicted protein sequence covered by matched peptides. MW (kDa): theoretical molecular mass calculated from the predicted amino-acid sequence. Peptide (Hits): number of unique tryptic peptides (and total spectra) matched to the protein.

Table S2. Table S2  Tryptic peptides identified by LC-MS/MS from the CCM1 immunoprecipitate and the 75 kDa (P75) and 45 kDa (P45) gel slices.

| | | | | | | P (pro) | Sf | Score | Coverage | MW | | Peptide (Hits) | Comments |
|---|---|---|---|---|---|---|---|---|---|---|---|---|---|
| Sample name | | Peptide | MH+ | DeltaM | z | P (pep) | Sf | XC | DeltaCn | Sp | RSp | Ions | |
| 01_CCM1 | 1 | estExt_fgenesh2_pg.C_50429 {CCM1} regulator of CO2-responsive genes | | | | 7.07E-13 | 7.50 | 80.30 | 21.30 | 70033.89 | | 9 (9 0 0 0 0) | |
| | | K.TLHGHDASLAVLPAPGGK.S | 1740.94 | 1.00 | 2 | 1.10E-08 | 0.97 | 4.21 | 0.54 | 1454.97 | 1.0 | 22/34 | |
| | | R.PGAALPPQAAPGTGAGQGAGAPAGAADGGAAPAAGDAAASGGAK.P | 3479.69 | 1.76 | 3 | 7.07E-13 | 0.95 | 4.78 | 0.55 | 1408.47 | 1.0 | 47/172 | |
| | | R.TGADDADEAVPGS*PHSK.H | 1733.70 | 1.06 | 2 | 1.97E-06 | 0.88 | 3.89 | 0.18 | 762.46 | 1.0 | 28/48 | phosphorylated fragment |
| | | R.TGADDADEAVPGSPHSK.H | 1653.74 | 1.96 | 2 | 5.71E-08 | 0.95 | 4.03 | 0.50 | 753.75 | 1.0 | 22/32 | |
| | | R.VSWLPGQVNGFVPQLQPQR.Y | 2150.15 | 1.58 | 3 | 3.52E-08 | 0.89 | 2.97 | 0.30 | 1434.91 | 1.0 | 31/72 | |
| | | R.DHLLDAETFR.L | 1216.60 | 0.18 | 2 | 5.84E-07 | 0.93 | 2.81 | 0.37 | 1210.01 | 1.0 | 15/18 | |
| | | K.QEHPEWPPTVPQGVFVGHGDR.V | 2369.14 | 1.73 | 3 | 5.86E-07 | 0.95 | 4.80 | 0.46 | 1103.86 | 1.0 | 32/80 | |
| | | K.PVADEDNLGTVFDDVEEFTR.D | 2268.03 | 0.64 | 2 | 3.21E-12 | 0.98 | 5.91 | 0.56 | 2470.27 | 1.0 | 28/38 | |
| | | K.PVADEDNLGTVFDDVEEFTR.D | 2268.03 | 1.90 | 3 | 3.20E-07 | 0.96 | 4.85 | 0.48 | 1519.17 | 1.0 | 29/76 | |
| | 2 | estExt_gwp_1H.C_900015 {ADH1} | | | | 1.02E-08 | 0.94 | 10.17 | | 102168.10 | | 1 (1 0 0 0 0) | |
| | | R.APATDEALTELK.A | 1258.65 | 0.99 | 2 | 1.02E-08 | 0.94 | 3.50 | 0.44 | 931.93 | 1.0 | 17/22 | |
| 02_74k | 1 | RWI_chlre3.14.27.2.51 Nickel chaperone for hydrogenase or urease, putative | | | | 3.28E-13 | 14.43 | 150.32 | 34.10 | 64549.51 | | 15 (15 0 0 0 0) | |
| | | R.TAGEAGPSGSSAADAK.A | 1376.63 | 0.46 | 2 | 3.69E-11 | 0.96 | 3.75 | 0.56 | 1319.44 | 1.0 | 23/30 | |
| | | K.AAVEDEAAAR.T | 1002.49 | 1.12 | 2 | 1.82E-05 | 0.96 | 3.47 | 0.44 | 1426.02 | 1.0 | 16/18 | |
| | | K.GQAAEPGAAAGADAGAGPGSK.A | 1710.80 | 1.30 | 2 | 3.28E-13 | 0.98 | 6.19 | 0.58 | 1728.52 | 1.0 | 32/40 | |
| | | K.IDLVNSEEQK.K | 1174.59 | 0.98 | 2 | 1.78E-06 | 0.95 | 3.56 | 0.44 | 1050.29 | 1.0 | 16/18 | |
| | | K.GVLSVHGYGSTK.F | 1204.63 | 0.76 | 2 | 1.87E-07 | 0.95 | 3.17 | 0.58 | 1132.18 | 1.0 | 18/22 | |
| | | K.IDLVNSEEQKK.Q | 1302.69 | 1.13 | 2 | 9.53E-07 | 0.95 | 3.71 | 0.41 | 912.46 | 1.0 | 17/20 | |
| | | R.YLMQHLEDGK.H | 1233.59 | 0.60 | 2 | 5.22E-07 | 0.96 | 3.23 | 0.56 | 1289.49 | 1.0 | 16/18 | |
| | | R.SIAVIENEFGEVNIDR.E | 1804.91 | 1.01 | 2 | 2.76E-11 | 0.98 | 5.34 | 0.51 | 2927.35 | 1.0 | 25/30 | |
| | | K.TQQPSQLLPR.R | 1167.65 | 0.50 | 2 | 1.83E-05 | 0.93 | 3.04 | 0.42 | 817.40 | 1.0 | 15/18 | |
| | | R.VNQVVFIGR.G | 1031.60 | 1.02 | 2 | 9.95E-06 | 0.96 | 3.85 | 0.38 | 1460.08 | 1.0 | 15/16 | |
| | | R.LHDLSDVSSVGIMAR.G | 1599.82 | 1.15 | 2 | 1.64E-12 | 0.98 | 5.21 | 0.62 | 2366.77 | 1.0 | 24/28 | |
| | | R.ELVAANLLAK.E | 1041.63 | 0.58 | 2 | 2.12E-05 | 0.93 | 2.43 | 0.41 | 1200.07 | 1.0 | 16/18 | |
| | | R.TASTVGQLALQQAAAAAASGAVAPATASAK.A | 2654.41 | 1.24 | 2 | 6.76E-11 | 0.98 | 6.47 | 0.56 | 1802.05 | 1.0 | 29/58 | |
| | | K.HSDGTVNEAVQQIAFADLILLNK.I | 2496.31 | 1.21 | 3 | 5.92E-06 | 0.96 | 4.59 | 0.52 | 1387.03 | 1.0 | 37/88 | |
| | | K.TPVTIITGFLGAGK.T | 1374.80 | 1.22 | 2 | 3.00E-10 | 0.97 | 4.49 | 0.54 | 1762.81 | 1.0 | 21/26 | |
| | 2 | estExt_fgenesh2_pg.C_50429 {CCM1} regulator of CO2-responsive genes | | | | 6.05E-12 | 0.98 | 10.29 | | 70033.89 | | 2 (2 0 0 0 0) | carry-over from the previous analysis |
| | | K.PVADEDNLGTVFDDVEEFTR.D | 2268.03 | 0.81 | 2 | 6.05E-12 | 0.98 | 5.89 | 0.56 | 2406.99 | 1.0 | 29/38 | |
| | | K.PVADEDNLGTVFDDVEEFTR.D | 2268.03 | 1.06 | 3 | 2.27E-05 | 0.92 | 3.86 | 0.40 | 1276.99 | 1.0 | 27/76 | |
| | 3 | Chlre2_kg.scaffold_1000769 | | | | 5.86E-07 | 0.96 | 10.17 | | 75461.40 | | 1 (1 0 0 0 0) | |
| | | R.SAVEGAEAGEGNGAGR.S | 1431.65 | 1.46 | 2 | 5.86E-07 | 0.96 | 3.40 | 0.42 | 1935.81 | 1.0 | 23/30 | |
| | 4 | Chlre2_kg.scaffold_1000770 | | | | 1.66E-07 | 0.95 | 10.16 | | 67702.49 | | 1 (1 0 0 0 0) | |
| | | K.LVGFDTSEPTAR.N | 1292.65 | 1.10 | 2 | 1.66E-07 | 0.95 | 3.22 | 0.49 | 1182.64 | 1.0 | 17/22 | |
| 03_45k | 1 | estExt_GenewiseW_1.C_760022 {GDH2} glutamate dehydrogenase | | | | 4.76E-10 | 3.75 | 40.23 | 9.30 | 49004.55 | | 4 (4 0 0 0 0) | |
| | | K.WEEDDVNR.K | 1062.45 | 1.32 | 2 | 1.68E-05 | 0.92 | 2.72 | 0.43 | 705.23 | 1.0 | 13/14 | |
| | | R.YHPQVDLDDVR.S | 1356.65 | 1.21 | 2 | 5.62E-07 | 0.89 | 2.85 | 0.32 | 612.33 | 1.0 | 16/20 | |
| | | K.PVYLHGSLGR.E | 1098.61 | 0.97 | 2 | 1.27E-07 | 0.96 | 3.09 | 0.50 | 1316.45 | 1.0 | 17/18 | |
| | | K.PLAEFTGGAAVPK.Q | 1257.68 | 1.23 | 2 | 4.76E-10 | 0.98 | 4.60 | 0.57 | 1695.21 | 1.0 | 21/24 | |
| | 2 | estExt_fgenesh1_pm.C_440001 {IDA5} Actin | | | | 6.30E-06 | 1.89 | 20.19 | | 41808.93 | | 2 (2 0 0 0 0) | |
| | | K.DSYVGDEAQSK.R | 1198.52 | 0.92 | 2 | 6.30E-06 | 0.94 | 2.91 | 0.46 | 1053.98 | 1.0 | 17/20 | |
| | | R.GYSFTTTAER.E | 1132.53 | 0.86 | 2 | 1.48E-05 | 0.95 | 3.04 | 0.53 | 907.78 | 1.0 | 16/18 | |
| | 3 | fgenesh2_pg.C_scaffold_2000011 | | | | 2.58E-05 | 0.93 | 10.18 | | 44510.41 | | 1 (1 0 0 0 0) | |
| | | K.EYVVQEGDVLLFR.F | 1566.82 | 1.47 | 2 | 2.58E-05 | 0.93 | 3.58 | 0.43 | 732.49 | 1.0 | 17/24 | |

Each band observed in Fig.1 was excised, digested with trypsin, and analysed by LC-MS/MS on a Q-TOF (Waters) or LTQ ion-trap (Thermo Scientific). For every sample the table lists all peptide-spectrum matches that passed the significance thresholds of Mascot (NAIST) or BioWorks (Okayama Univ.). Peptides in which phosphorylation was detected are highlighted in blue.

Columns. Peptide: identified peptide sequence (asterisk denotes a post-translational modification, here phosphorylation). $MH^+$: monoisotopic mass of the protonated peptide (Da). ΔM: mass error relative to the theoretical value (Da). z: precursor ion charge state. P (pro)/P (pep) and Sf: Mascot or BioWorks probability and score for the corresponding protein (P (pro)) or individual peptide (P (pep)). XC: cross-correlation score. ΔCn: normalised difference between the best and second-best matches. Sp and RSp: preliminary score and rank of preliminary score. Coverage and MW for each protein are reproduced from Table S1. Ions: number of matched fragment ions (y + b) over total considered.

Table S3. Proteins co-purifying with CCM1-FLAG under high- and very-low-$CO_2$ conditions, identified by in-solution tryptic digestion.

| Reference | P (pro) | Sf | Score | Coverage | MW | Peptide (Hits) |
|---|---|---|---|---|---|---|
| High-CO2 conditions | | | | | | |
| jgi\|Chlre3\|195946\|RWI_chlre3.14.27.2.51 Nickel chaperone for hydrogenase or urease, putative | 3.33E-15 | 18.96 | 200.35 | 45.90 | 64549.51 | 27 (27 0 0 0 0) |
| jgi\|Chlre3\|82916\|estExt_GenewiseW_1.C_760022 {GDH2} glutamate dehydrogenase | 1.72E-13 | 8.46 | 90.27 | 24.90 | 49004.55 | 9 (9 0 0 0 0) |
| jgi\|Chlre3\|186972\|estExt_fgenesh2_pg.C_50429 {CCM1} regulator of CO2-responsive genes | 6.27E-13 | 7.58 | 80.29 | 25.50 | 70033.89 | 8 (8 0 0 0 0) |
| jgi\|Chlre3\|190969\|estExt_fgenesh2_pg.C_220037 {GDH1} | 1.74E-08 | 5.69 | 60.25 | 19.00 | 48534.99 | 6 (6 0 0 0 0) |
| jgi\|Chlre3\|134382\|estExt_gwp_1W.C_30492 {EF1A1} eukaryotic translation elongation factor 1 alpha 1 | 1.17E-09 | 3.64 | 40.19 | 11.40 | 50792.30 | 4 (4 0 0 0 0) |
| jgi\|Chlre3\|137452\|estExt_gwp_1W.C_220126 {HSP70C} Heat shock protein 70C | 1.22E-12 | 2.91 | 30.25 | 8.70 | 65263.79 | 3 (3 0 0 0 0) |
| jgi\|Chlre3\|188942\|estExt_fgenesh2_pg.C_120244 {IF4A} Similar to Eukaryotic Initiation Factor 4A | 5.03E-12 | 1.94 | 20.21 | 7.00 | 47024.02 | 2 (2 0 0 0 0) |
| jgi\|Chlre3\|128745\|estExt_gwp_1H.C_150086 {RCA} rubisco activase | 2.04E-13 | 1.87 | 20.25 | 10.80 | 45000.80 | 2 (2 0 0 0 0) |
| jgi\|Chlre3\|10441\|fgenesh1_pg.C_scaffold_11000230 {FBP} fructose-1,6-bisphosphatase | 6.56E-07 | 1.87 | 20.19 | 8.00 | 35372.45 | 2 (2 0 0 0 0) |
| Reference | P (pro) | Sf | Score | Coverage | MW | Peptide (Hits) |
| Very low-CO2 conditions | | | | | | |
| jgi\|Chlre3\|195946\|RWI_chlre3.14.27.2.51 Nickel chaperone for hydrogenase or urease, putative | 1.67E-14 | 18.87 | 200.35 | 43.20 | 64549.5 | 23 (23 0 0 0 0) |
| jgi\|Chlre3\|82916\|estExt_GenewiseW_1.C_760022 {GDH2} glutamate dehydrogenase | 1.43E-11 | 11.33 | 120.29 | 34.90 | 49004.6 | 12 (12 0 0 0 0) |
| jgi\|Chlre3\|186972\|estExt_fgenesh2_pg.C_50429 {CCM1} regulator of CO2-responsive genes | 1.11E-16 | 10.51 | 110.34 | 22.90 | 70033.9 | 12 (12 0 0 0 0) |
| jgi\|Chlre3\|190969\|estExt_fgenesh2_pg.C_220037 {GDH1} | 1.79E-12 | 7.69 | 80.31 | 28.80 | 48535.0 | 8 (8 0 0 0 0) |
| jgi\|Chlre3\|191668\|estExt_fgenesh2_pg.C_260094 {ICL1} isocitrate lyase | 9.48E-12 | 3.79 | 40.25 | 19.70 | 45719.7 | 4 (4 0 0 0 0) |
| jgi\|Chlre3\|137452\|estExt_gwp_1W.C_220126 {HSP70C} Heat shock protein 70C | 1.38E-11 | 2.94 | 30.29 | 8.90 | 65263.8 | 3 (3 0 0 0 0) |
| jgi\|Chlre3\|104082\|e_gwH.26.231.1 | 3.05E-09 | 3.89 | 40.28 | 65.50 | 8961.9 | 4 (4 0 0 0 0) |
| jgi\|Chlre3\|183343\|estExt_fgenesh2_kg.C_140079 | 1.07E-11 | 2.83 | 30.24 | 15.30 | 25643.5 | 3 (3 0 0 0 0) |
| jgi\|Chlre3\|144137\|Chlre2_kg.scaffold_10000017 | 4.93E-06 | 1.73 | 20.16 | 14.90 | 15907.2 | 2 (2 0 0 0 0) |

FLAG-tagged CCM1 complexes were affinity-purified from cells cultured continuously at 5 % $CO_2$ (high $CO_2$, HC) or from cells shifted for 12 h to 0.04 % $CO_2$ (very-low $CO_2$, VLC). Eluates were reduced, alkylated, and digested with trypsin in solution before LC-MS/MS analysis on an LTQ ion-trap (Thermo Scientific) at Okayama University. Spectra were searched with BioWorks against the Chlamydomonas reinhardtii database (JGI v3.1). Only proteins represented by ≥ 2 unique peptides in at least two technical replicates for either condition are reported. Columns are as follows. Reference: JGI protein ID and annotation returned as the top hit. P (pro): ranking of the hit within the sample. Sf: Mascot/BioWorks scoring function (lower E-value or higher score indicates greater confidence). Score: confidence score assigned by the search programme. Coverage (%): percentage of the predicted protein sequence covered by matched peptides. MW (kDa): theoretical molecular mass calculated from the predicted amino-acid sequence. Peptide (Hits): number of unique tryptic peptides (and total spectra) matched to the protein. Proteins that were already identified by the earlier in-gel digestion workflow (Table S1 and S2) are highlighted in yellow to facilitate comparison between the two methodologies.

Table S4.  Peptide map of the excised CCM1-FLAG band and its phosphorylation sites.

| | | | | | | P (pro) | Sf | Score | Coverage | MW | | Peptide (Hits) | Comments |
|---|---|---|---|---|---|---|---|---|---|---|---|---|---|
| Sample name | | Peptide | MH+ | DeltaM | z | P (pep) | Sf | XC | DeltaCn | Sp | RSp | Ions | |
| CCM1-HC | 1 | estExt_fgenesh2_pg.C_50429 {CCM1} regulator of CO2-responsive genes | | | | | | | | | | | |
| | | -.M]EALDAQDSLQLDVVSPSAR.P | 2187.06 | 0.46 | 2 | 8.87E-11 | 0.95 | 3.57 | 0.54 | 977.3 | 1 | 24/38 | |
| | | K.PVADEDNLGTVFDDVEEFTR.D | 2268.03 | 0.86 | 2 | 6.67E-11 | 0.98 | 6.19 | 0.56 | 1972.4 | 1 | 26/38 | |
| | | K.SWQDFALTHAGYAIGAPAMLAPLK.Q | 2529.30 | 1.02 | 3 | 1.66E-04 | 0.88 | 3.87 | 0.29 | 982.2 | 1 | 31/92 | |
| | | K.TLHGHDASLAVLPAPGGK.S | 1740.94 | 1.17 | 2 | 8.31E-08 | 0.95 | 3.78 | 0.49 | 1213.4 | 1 | 20/34 | |
| | | R.DHLLDAETFR.L | 1216.60 | 0.58 | 2 | 2.03E-05 | 0.93 | 2.98 | 0.34 | 1228.6 | 1 | 15/18 | |
| | | R.IPS*PPPLPPDFHTAATGGNGMLFNFSQFGQK.L | 3350.57 | 0.75 | 3 | 9.46E-12 | 0.94 | 5.77 | 0.25 | 999.5 | 1 | 49/180 | Phosphorylated |
| | | R.PGAALPPQAAPGTGAGQGAGAPAGAADGGAAPAAGDAAASGGAK.P | 3479.69 | 1.39 | 3 | 2.39E-11 | 0.97 | 5.53 | 0.65 | 1499.4 | 1 | 48/172 | |
| | | R.TGADDADEAVPGS*PHSK.H | 1733.70 | 0.98 | 2 | 5.99E-06 | 0.89 | 4.08 | 0.18 | 802.9 | 1 | 27/48 | Phosphorylated |
| | | R.VSWLPGQVNGFVPQLQPQR.Y | 2150.15 | 1.20 | 2 | 3.15E-10 | 0.98 | 4.91 | 0.46 | 1972.9 | 1 | 27/36 | |
| CCM1-VLC | 1 | estExt_fgenesh2_pg.C_50429 {CCM1} regulator of CO2-responsive genes | | | | | | | | | | | |
| | | -.M]EALDAQDSLQLDVVS*PSARPAAAGGDKR.D | 3090.46 | 1.19 | 3 | 1.71E-10 | 0.96 | 5.45 | 0.09 | 2499.9 | 1 | 57/168 | Phosphorylated |
| | | -.M]EALDAQDSLQLDVVSPSAR.P | 2187.06 | 1.46 | 2 | 1.17E-11 | 0.97 | 4.00 | 0.59 | 1411.8 | 1 | 28/38 | |
| | | -.M]EALDAQDSLQLDVVSPSARPAAAGGDKR.D | 3010.49 | 1.81 | 3 | 2.22E-15 | 0.97 | 5.86 | 0.53 | 1361.4 | 1 | 39/112 | |
| | | K.PVADEDNLGTVFDDVEEFTR.D | 2268.03 | 0.99 | 2 | 3.96E-12 | 0.99 | 6.19 | 0.58 | 2364.8 | 1 | 28/38 | |
| | | K.SLMNGGAGHAGEEHHR.D | 1659.74 | 1.28 | 2 | 7.02E-09 | 0.94 | 4.02 | 0.52 | 887.8 | 1 | 18/30 | |
| | | K.SLMNGGAGHAGEEHHRDHLLDAETFR.L | 2857.32 | 1.34 | 3 | 1.31E-07 | 0.88 | 3.89 | 0.35 | 938.9 | 1 | 29/100 | |
| | | K.TLHGHDASLAVLPAPGGK.S | 1740.94 | -0.24 | 2 | 1.31E-07 | 0.96 | 3.85 | 0.49 | 1438.1 | 1 | 22/34 | |
| | | R.DHLLDAETFR.L | 1216.60 | 0.59 | 2 | 6.90E-05 | 0.95 | 2.95 | 0.38 | 1369.6 | 1 | 16/18 | |
| | | R.IPS*PPPLPPDFHTAATGGNGMLFNFSQFGQK.L | 3350.57 | -0.40 | 3 | 7.05E-13 | 0.97 | 6.49 | 0.35 | 1703.0 | 1 | 52/180 | Phosphorylated |
| | | R.PGAALPPQAAPGTGAGQGAGAPAGAADGGAAPAAGDAAASGGAK.P | 3479.69 | 0.75 | 3 | 2.11E-14 | 0.97 | 5.56 | 0.62 | 1994.2 | 1 | 53/172 | |
| | | R.SFAELWR.L | 908.46 | 0.97 | 2 | 1.86E-04 | 0.88 | 2.10 | 0.42 | 592.4 | 1 | 11/12 | |
| | | R.TGADDADEAVPGS*PHSK.H | 1733.70 | 0.98 | 2 | 5.99E-06 | 0.89 | 4.08 | 0.18 | 802.9 | 1 | 27/48 | Phosphorylated |
| | | R.TGADDADEAVPGSPHSK.H | 1653.74 | 0.90 | 2 | 1.11E-07 | 0.93 | 3.83 | 0.48 | 617.0 | 1 | 21/32 | |
| | | R.VSWLPGQVNGFVPQLQPQR.Y | 2150.15 | 0.74 | 2 | 1.19E-12 | 0.98 | 5.44 | 0.47 | 1913.3 | 1 | 27/36 | |

The single CCM1-FLAG band was cut from an SDS-PAGE gel after FLAG affinity purification, reduced, alkylated, and subjected to in-gel tryptic digestion. Resulting peptides were analysed by LC-MS/MS on an LTQ ion-trap (Thermo Scientific) at Okayama University, and spectra were searched with BioWorks against the Chlamydomonas reinhardtii database (JGI v3.1). The table lists every CCM1 peptide that met the BioWorks significance threshold in at least two technical replicates of either $CO_2$ treatment—continuous high $CO_2$ (5 %, HC) or a 12 h shift to very-low $CO_2$ (0.04 %, VLC). Peptides that contain a phosphorylation site are shaded light blue.

Table S5. Photosynthetic parameters of WT and transformant cells

| Growth condition | Measuring condition | Strain name | Vmax of $O_2$-evolving activity [μmol $O_2$·mgChl$^{-1}$·h$^{-1}$] | $K_{0.5}$(Ci) [μM] |
|---|---|---|---|---|
| VLC | 20 mM HEPES-NaOH (pH 7.8) | WT | 229 ± 40 | 28 ± 4 |
| VLC | 20 mM HEPES-NaOH (pH 7.8) | *cbp1*-1 | 221 ± 26 | 29 ± 5 |
| VLC | 20 mM HEPES-NaOH (pH 7.8) | *cbp1*-1:*CBP1* | 219 ± 15 | 20 ± 5 |
| VLC | 20 mM HEPES-NaOH (pH 7.8) | *ccm1*-1 | 224 ± 52 | 254 ± 111 |
| VLC | 20 mM HEPES-NaOH (pH 7.8) | *ccm1*-1:*CCM1* | 217 ± 10 | 22 ± 4 |
| HC | 20 mM HEPES-NaOH (pH 7.8) | WT | 212 ± 32 | 381 ± 26 |
| HC | 20 mM HEPES-NaOH (pH 7.8) | *cbp1*-1 | 178 ± 11 | 248 ± 27 |
| HC | 20 mM HEPES-NaOH (pH 7.8) | *cbp1*-1:*CBP1* | 202 ± 9 | 475 ± 40 |
| HC | 20 mM HEPES-NaOH (pH 7.8) +1% [v/v] DMSO | WT | 208 ± 11 | 394 ± 31 |
| HC | 20 mM HEPES-NaOH (pH 7.8) +1% [v/v] DMSO | *cbp1*-1 | 203 ± 23 | 169 ± 5 |
| HC | 20 mM HEPES-NaOH (pH 7.8) + 5 mM AZA in DMSO | *cbp1*-1 | 213 ± 38 | 416 ± 70 |

The data are shown ± standard error (SE), which were obtained from three independent experiments. AZA, acetazolamide; DMSO, Dimethyl sulfoxide; HC, high-$CO_2$; $K_{0.5}$(Ci), Ci concentration required for half of Vmax; VLC, very low-$CO_2$; Vmax, maximum $O_2$-evolving activity.

Table S6. VLC-inducible and CCM1-dependent genes differently expressed in *cbp1*-1 cells compared with those in WT and *cbp1*-1:*CBP1* cells in HC conditions

| Gene ID | Gene name | Description | Average TPM in WT | | | Average TPM in *ccm1*-1 | | | Average TPM in *ccm1*-1:*CCM1* | | | Average TPM in *cbp1*-1 | | | Average TPM in *cbp1*-1:*CBP1* | | | WT_HC vs *cbp1*-1_HC | | *cbp1*-1:*CBP1*_HC vs *cbp1*-1_HC | | WT_VLC 0.3 h vs *cbp1*-1_VLC 0.3 h | | *cbp1*-1:*CBP1*_VLC 0.3 h vs *cbp1*-1_VLC 0.3 h | | WT_VLC 2.0 h vs *cbp1*-1_VLC 2.0 h | | *cbp1*-1:*CBP1*_VLC 2.0 h vs *cbp1*-1_VLC 2.0 h | |
|---|---|---|---|---|---|---|---|---|---|---|---|---|---|---|---|---|---|---|---|---|---|---|---|---|---|---|---|---|---|
| | | | HC | VLC 0.3 h | VLC 2.0 h | HC | VLC 0.3 h | VLC 2.0 h | HC | VLC 0.3 h | VLC 2.0 h | HC | VLC 0.3 h | VLC 2.0 h | HC | VLC 0.3 h | VLC 2.0 h | $log_2FC$ | FDR | $log_2FC$ | FDR | $log_2FC$ | FDR | $log_2FC$ | FDR | $log_2FC$ | FDR | $log_2FC$ | FDR |
| **Cre01.g000150** | **ZRT2** | Zinc-nutrition responsive permease transporter | 3.8 | 32.0 | 8.8 | 1.8 | 0.9 | 0.4 | 4.0 | 24.2 | 11.0 | 13.0 | 30.2 | 8.8 | 4.1 | 27.0 | 8.2 | 1.34 | 2.2.E-04 | 1.78 | 1.1.E-13 | -0.40 | 4.9.E-01 | 0.42 | 3.0.E-01 | -0.45 | 5.4.E-01 | 0.01 | 1.0.E+00 |
| **Cre01.g053950** | — | Factors identified as interacting with LCIB/LCIC in the proteome analysis by Mackinder et al. Cell (2017) | 0.7 | 50.2 | 28.7 | 0.1 | 0.2 | 0.5 | 1.6 | 37.7 | 34.0 | 12.3 | 80.3 | 48.6 | 3.2 | 84.8 | 36.9 | 3.65 | 2.0.E-34 | 2.05 | 1.9.E-19 | 0.36 | 4.9.E-01 | 0.17 | 7.4.E-01 | 0.29 | 7.0.E-01 | 0.30 | 8.6.E-01 |
| **Cre03.g151650** | ***SMM7*** | Putative methyltransferase that localizes to the pyrenoid matrix and is implicated in pyrenoid biogenesis (Mackinder et al. Cell 2017) | 7.7 | 252.3 | 107.6 | 0.1 | 0.2 | 0.1 | 13.5 | 217.1 | 91.9 | 27.0 | 144.8 | 47.9 | 8.5 | 163.7 | 51.9 | 1.37 | 1.3.E-03 | 1.78 | 2.7.E-13 | -1.09 | 3.6.E-03 | 0.09 | 8.5.E-01 | -1.62 | 3.3.E-05 | -0.21 | 9.3.E-01 |
| **Cre03.g162800** | ***LCI1*** | Low-CO2-inducible membrane protein | 0.1 | 1270.3 | 947.6 | 0.0 | 0.5 | 0.1 | 0.2 | 991.2 | 1116.6 | 1.0 | 1490.1 | 709.4 | 0.1 | 456.1 | 307.1 | 2.36 | 3.6.E-03 | 3.75 | 6.7.E-05 | -0.12 | 8.8.E-01 | 1.94 | 1.1.E-10 | -0.91 | 2.6.E-01 | 1.12 | 1.2.E-01 |
| **Cre03.g212641** | — | Putative histone acetyltransferase of the CBP family 1 (Li et al. Genomics 2020) | 1.9 | 14.4 | 3.2 | 0.5 | 1.1 | 1.5 | 1.3 | 10.1 | 4.4 | 12.5 | 17.0 | 10.6 | 5.0 | 20.5 | 8.8 | 2.30 | 1.3.E-14 | 1.44 | 2.1.E-07 | -0.08 | 8.9.E-01 | -0.03 | 9.6.E-01 | 1.27 | 1.0.E-02 | 0.18 | 9.5.E-01 |
| **Cre03.g212977** | — | Putative histone acetyltransferase of the CBP family 1 (Li et al. Genomics 2020) | 1.1 | 40.0 | 31.6 | 0.0 | 0.0 | 0.0 | 1.7 | 25.7 | 45.3 | 18.0 | 54.2 | 38.0 | 5.7 | 43.6 | 28.3 | 3.58 | 3.4.E-26 | 1.78 | 2.8.E-12 | 0.10 | 8.8.E-01 | 0.54 | 2.7.E-01 | -0.20 | 8.1.E-01 | 0.33 | 8.9.E-01 |
| **Cre04.g215900** | — | Tyrosine-protein kinase ephrin type A/B receptor-like protein | 0.4 | 14.1 | 6.1 | 0.2 | 0.1 | 0.0 | 0.5 | 9.6 | 6.7 | 7.8 | 21.3 | 11.5 | 3.6 | 19.5 | 10.3 | 3.90 | 1.5.E-22 | 1.24 | 1.1.E-05 | 0.23 | 7.6.E-01 | 0.35 | 4.9.E-01 | 0.45 | 4.7.E-01 | 0.06 | 9.9.E-01 |
| **Cre04.g215952** | — | — | 0.3 | 2.4 | 2.3 | 0.1 | 0.0 | 0.1 | 0.1 | 1.9 | 1.4 | 11.0 | 37.4 | 21.2 | 4.3 | 28.3 | 16.3 | 4.69 | 1.4.E-20 | 1.44 | 1.6.E-04 | 3.67 | 1.2.E-19 | 0.68 | 3.5.E-02 | 2.71 | 5.2.E-10 | 0.28 | 9.0.E-01 |
| **Cre04.g222750** | ***CCP2*** | Low-CO2-inducible membrane protein | 3.8 | 226.0 | 235.4 | 2.0 | 4.1 | 2.7 | 8.1 | 131.3 | 318.9 | 26.2 | 529.9 | 314.0 | 12.5 | 266.7 | 234.0 | 2.32 | 2.2.E-12 | 1.17 | 6.0.E-06 | 0.87 | 2.3.E-01 | 1.23 | 2.9.E-04 | -0.09 | 9.5.E-01 | 0.33 | 9.4.E-01 |
| **Cre04.g222800** | ***LCID*** | Low-CO2 inducible protein D | 2.8 | 158.5 | 236.4 | 2.1 | 3.2 | 3.4 | 3.2 | 101.2 | 324.7 | 10.5 | 253.8 | 174.2 | 5.3 | 141.5 | 105.5 | 1.44 | 3.2.E-06 | 1.08 | 1.4.E-04 | 0.35 | 5.5.E-01 | 1.09 | 7.6.E-03 | -0.91 | 1.1.E-01 | 0.63 | 6.3.E-01 |
| **Cre04.g223100** | ***CAH1*** | Periplasmic carbonic anhydrase | 13.9 | 5174.3 | 8334.0 | 2.4 | 4.2 | 2.7 | 19.0 | 5047.9 | 9969.0 | 114.1 | 2696.0 | 8316.8 | 3.1 | 1751.1 | 5037.5 | 2.60 | 5.6.E-22 | 5.32 | 1.5.E-75 | -1.23 | 1.4.E-03 | 0.88 | 6.3.E-03 | -0.43 | 5.5.E-01 | 0.63 | 6.9.E-01 |
| **Cre06.g278199** | ***LOCO2*** | Low CO2 sensitive 2 (Fauser et al. Nat Genet 2022) | 0.9 | 3.1 | 4.0 | 1.4 | 2.3 | 1.2 | 2.0 | 5.9 | 7.9 | 9.0 | 29.5 | 20.9 | 4.5 | 23.5 | 10.3 | 2.92 | 1.5.E-16 | 1.08 | 4.7.E-04 | 2.97 | 1.1.E-15 | 0.60 | 5.4.E-02 | 1.92 | 2.5.E-06 | 0.93 | 2.6.E-02 |
| **Cre06.g309000** | ***LCIA*** | Inorganic carbon channel localized at chloroplast membrane | 0.6 | 1456.6 | 1270.7 | 0.0 | 0.6 | 0.0 | 0.4 | 990.5 | 1588.1 | 5.0 | 2114.9 | 1347.6 | 0.4 | 1189.6 | 813.5 | 2.65 | 2.6.E-07 | 3.66 | 7.4.E-12 | 0.21 | 7.6.E-01 | 1.08 | 4.9.E-03 | -0.39 | 6.5.E-01 | 0.64 | 7.4.E-01 |
| **Cre07.g339000** | ***RbcX-Iib*** | Chaperon-like RbcX protein (Bracher et al. PLOS One 2015) | 1.0 | 18.0 | 11.4 | 0.7 | 0.5 | 0.4 | 2.5 | 14.3 | 12.7 | 8.3 | 29.5 | 16.1 | 3.8 | 23.8 | 12.4 | 2.53 | 9.2.E-09 | 1.23 | 4.0.E-03 | 0.45 | 3.4.E-01 | 0.59 | 8.6.E-02 | 0.04 | 9.6.E-01 | 0.28 | 8.6.E-01 |
| **Cre08.g367400** | ***LHCSR3.2*** | Stress-related chlorophyll a/b binding protein 3 | 0.3 | 824.0 | 684.3 | 0.1 | 4.4 | 3.1 | 2.0 | 635.7 | 512.4 | 6.3 | 1569.0 | 686.7 | 1.9 | 957.0 | 399.7 | 3.86 | 5.1.E-15 | 1.86 | 1.2.E-06 | 0.62 | 2.1.E-01 | 0.98 | 4.5.E-03 | -0.45 | 4.7.E-01 | 0.69 | 8.0.E-01 |
| **Cre08.g367500** | ***LHCSR3.1*** | Stress-related chlorophyll a/b binding protein 2 | 0.6 | 655.5 | 582.2 | 0.3 | 2.4 | 2.2 | 1.3 | 477.1 | 442.2 | 5.2 | 1354.3 | 616.3 | 1.7 | 782.6 | 337.7 | 2.70 | 3.8.E-07 | 1.73 | 4.4.E-05 | 0.74 | 1.6.E-01 | 1.06 | 6.2.E-04 | -0.38 | 5.8.E-01 | 0.78 | 7.2.E-01 |
| **Cre09.g394473** | ***LCI9*** | Low-CO2 inducible protein 9 containing starch-binding domain of CBM_20 (pfam00686) | 14.7 | 353.4 | 240.4 | 0.5 | 0.6 | 1.6 | 21.2 | 306.7 | 271.1 | 61.0 | 229.0 | 151.5 | 18.3 | 300.5 | 164.3 | 1.61 | 4.0.E-06 | 1.84 | 5.2.E-12 | -0.93 | 1.9.E-02 | -0.15 | 7.6.E-01 | -1.10 | 8.7.E-03 | -0.21 | 9.6.E-01 |
| **Cre09.g395732** | — | DnaJ domain containing protein | 0.8 | 17.6 | 2.1 | 0.6 | 1.9 | 3.1 | 0.7 | 13.9 | 4.3 | 4.4 | 21.7 | 5.2 | 1.5 | 28.4 | 5.5 | 1.98 | 3.0.E-09 | 1.62 | 8.5.E-09 | -0.03 | 9.6.E-01 | -0.14 | 8.2.E-01 | 0.81 | 1.7.E-01 | -0.19 | 9.5.E-01 |
| **Cre09.g399552** | ***LCR1*** | Low-CO2 response regulator, Myb-like transcription factor | 0.5 | 54.6 | 53.2 | 0.1 | 0.9 | 1.1 | 0.5 | 22.6 | 60.7 | 13.2 | 61.3 | 74.8 | 1.0 | 37.6 | 44.5 | 4.20 | 4.5.E-33 | 3.81 | 8.8.E-39 | -0.19 | 7.9.E-01 | 0.93 | 4.1.E-02 | 0.02 | 9.8.E-01 | 0.66 | 2.9.E-01 |
| **Cre10.g452800** | ***LCIB*** | Low-CO2-inducible protein B | 172.0 | 5242.3 | 901.0 | 58.1 | 181.8 | 187.7 | 202.6 | 5126.6 | 1390.4 | 612.7 | 4717.7 | 1372.4 | 149.3 | 4493.7 | 1281.3 | 1.39 | 6.4.E-07 | 2.14 | 1.0.E-15 | -0.45 | 3.5.E-01 | 0.33 | 4.4.E-01 | 0.16 | 8.4.E-01 | 0.00 | 1.0.E+00 |
| **Cre11.g481104** | — | protein kinase family protein | 1.6 | 22.4 | 8.7 | 1.6 | 2.0 | 1.5 | 1.3 | 16.3 | 12.6 | 6.2 | 24.8 | 10.9 | 3.3 | 25.2 | 8.1 | 1.50 | 1.2.E-04 | 1.02 | 2.1.E-03 | -0.17 | 7.7.E-01 | 0.23 | 6.8.E-01 | -0.13 | 8.6.E-01 | 0.33 | 7.7.E-01 |
| **Cre16.g662600** | ***BST1*** | Thylakoid localized bestrophin-like protein 1 | 0.1 | 116.6 | 8.3 | 0.0 | 0.1 | 0.0 | 0.1 | 55.0 | 10.2 | 11.7 | 183.5 | 31.2 | 0.4 | 155.1 | 25.8 | 6.39 | 1.5.E-38 | 4.93 | 7.5.E-42 | 0.28 | 7.6.E-01 | 0.46 | 3.9.E-01 | 1.42 | 2.5.E-02 | 0.18 | 9.6.E-01 |
| **Cre16.g663450** | ***BST3/LCI11*** | Thylakoid localized bestrophin-like protein 3 | 40.9 | 700.5 | 359.2 | 16.8 | 22.2 | 17.6 | 43.0 | 679.9 | 455.5 | 125.0 | 826.1 | 346.1 | 41.4 | 673.4 | 233.3 | 1.15 | 3.8.E-03 | 1.70 | 1.6.E-05 | -0.04 | 9.4.E-01 | 0.56 | 9.8.E-02 | -0.50 | 3.8.E-01 | 0.47 | 6.6.E-01 |
| **Cre16.g667250** | — | — | 26.9 | 194.5 | 194.6 | 22.7 | 20.9 | 48.9 | 34.6 | 173.9 | 252.4 | 87.8 | 358.2 | 252.8 | 46.4 | 258.3 | 215.3 | 1.26 | 8.4.E-05 | 1.02 | 4.2.E-03 | 0.60 | 2.0.E-01 | 0.75 | 2.8.E-02 | -0.08 | 9.2.E-01 | 0.14 | 9.9.E-01 |
| **Cre16.g674291** | — | Protein kinase family protein | 0.9 | 16.2 | 6.1 | 0.9 | 1.3 | 0.7 | 1.1 | 10.5 | 8.4 | 19.4 | 32.6 | 34.1 | 9.9 | 34.4 | 19.7 | 4.00 | 3.2.E-42 | 1.07 | 1.8.E-05 | 0.71 | 9.2.E-02 | 0.18 | 6.6.E-01 | 2.02 | 8.1.E-08 | 0.70 | 6.8.E-02 |
| **Cre16.g685050** | ***LCI15*** | Cobalamin synthesis protein cobW C-terminal domain (CobW_C) and WW domain containing protein | 17.9 | 225.5 | 153.1 | 8.9 | 20.0 | 16.9 | 13.6 | 175.7 | 243.3 | 71.5 | 324.9 | 314.5 | 19.3 | 257.4 | 203.3 | 1.54 | 5.0.E-07 | 1.99 | 8.2.E-15 | 0.23 | 6.3.E-01 | 0.60 | 6.0.E-02 | 0.58 | 2.9.E-01 | 0.54 | 5.4.E-01 |
| **Cre16.g685100** | — | Cobalamin synthesis protein cobW C-terminal domain (CobW_C) and WW domain containing protein | 0.5 | 11.6 | 7.9 | 0.0 | 0.0 | 0.1 | 0.3 | 4.4 | 7.7 | 1.8 | 22.1 | 17.9 | 0.4 | 9.2 | 9.9 | 1.52 | 2.5.E-03 | 2.37 | 3.0.E-06 | 0.57 | 3.5.E-01 | 1.48 | 2.3.E-04 | 0.67 | 4.4.E-01 | 0.76 | 4.3.E-01 |

Gene IDs represent the gene accession numbers from Phytozome.

Table S7. VLC-inducible and CCM1-dependent genes differently expressed in *cbp1*-1 cells compared with those in WT and *cbp1*-1:*CBP1* cells in VLC conditions

| Gene ID | Gene name | Description | Average TPM in WT | | | Average TPM in *ccm1*-1 | | | Average TPM in *ccm1*-1:*CCM1* | | | Average TPM in *cbp1*-1 | | | Average TPM in *cbp1*-1:*CBP1* | | |
|---|---|---|---|---|---|---|---|---|---|---|---|---|---|---|---|---|---|
| | | | HC | VLC 0.3 h | VLC 2.0 h | HC | VLC 0.3 h | VLC 2.0 h | HC | VLC 0.3 h | VLC 2.0 h | HC | VLC 0.3 h | VLC 2.0 h | HC | VLC 0.3 h | VLC 2.0 h |
| **Cre01.g047650** | — | F-box protein with leucine-rich repeats protein | 1.4 | 8.6 | 9.5 | 1.3 | 2.0 | 1.1 | 1.2 | 5.5 | 14.8 | 4.4 | 27.0 | 13.2 | 3.8 | 13.7 | 8.1 |
| **Cre03.g198250** | — | LCI12 paralog | 422.8 | 1134.5 | 515.9 | 104.6 | 136.2 | 163.9 | 321.1 | 800.4 | 438.6 | 178.2 | 226.5 | 82.5 | 160.0 | 384.8 | 196.3 |
| **Cre03.g204465** | — | GT90 family protein 36 | 0.2 | 1.2 | 5.0 | 0.1 | 0.1 | 0.1 | 0.2 | 0.5 | 4.4 | 1.3 | 6.5 | 9.3 | 2.0 | 2.8 | 6.2 |
| **Cre03.g204577** | ***DNJ31*** | DnaJ-like protein | 0.0 | 5.8 | 32.5 | 0.0 | 0.0 | 0.0 | 0.0 | 1.5 | 39.3 | 0.1 | 36.7 | 52.0 | 0.2 | 7.4 | 16.4 |
| **Cre04.g216550** | — | — | 0.9 | 4.7 | 5.5 | 1.2 | 8.9 | 7.5 | 0.8 | 5.2 | 7.6 | 2.1 | 14.3 | 15.4 | 1.1 | 7.3 | 6.9 |
| **Cre04.g223250** | ***LCIE*** | Low-CO2 inducible protein E | 0.3 | 7.4 | 6.4 | 0.1 | 0.0 | 0.0 | 0.3 | 2.7 | 8.2 | 0.4 | 49.8 | 33.1 | 0.1 | 9.2 | 12.0 |
| **Cre05.g245500** | ***FAP175*** | Ankyrin Repeat Flagellar Associated Protein 175 | 0.7 | 6.5 | 3.7 | 0.8 | 1.0 | 0.5 | 0.3 | 3.2 | 4.0 | 1.2 | 2.3 | 2.4 | 3.7 | 10.6 | 6.5 |
| **Cre08.g380700** | — | — | 2.7 | 5.1 | 17.0 | 5.1 | 4.5 | 5.6 | 6.1 | 6.0 | 19.9 | 13.8 | 22.1 | 33.8 | 8.6 | 11.5 | 18.7 |
| **Cre11.g469650** | — | — | 1.9 | 4.9 | 8.2 | 2.4 | 1.7 | 4.2 | 1.6 | 3.9 | 7.9 | 3.8 | 14.4 | 11.6 | 4.2 | 6.8 | 8.4 |
| **Cre12.g527250** | — | — | 0.8 | 5.2 | 6.6 | 0.9 | 1.1 | 0.7 | 0.2 | 1.7 | 4.6 | 2.0 | 23.9 | 22.7 | 3.9 | 9.2 | 12.6 |
| **Cre12.g555700** | ***DNJ15*** | DnaJ-like protein | 0.1 | 5.8 | 83.1 | 0.0 | 0.0 | 0.0 | 0.0 | 0.6 | 96.7 | 0.2 | 48.5 | 110.0 | 0.1 | 7.5 | 43.0 |
| **Cre16.g652200** | ***MMP9*** | Metalloproteinase of VMP family | 3.8 | 21.3 | 5.7 | 2.4 | 21.4 | 8.0 | 6.5 | 32.1 | 9.7 | 1.3 | 3.2 | 2.7 | 2.5 | 8.6 | 4.2 |
| **Cre17.g719150** | — | LCI12 paralog | 0.3 | 5.1 | 7.1 | 0.1 | 0.0 | 0.0 | 0.4 | 4.0 | 4.8 | 2.2 | 3.2 | 0.8 | 3.6 | 9.0 | 10.8 |
| **Cre19.g751047** | — | LCI12 paralog | 429.8 | 1526.0 | 857.7 | 71.1 | 92.8 | 123.9 | 351.0 | 1080.9 | 717.1 | 304.9 | 412.5 | 155.0 | 417.1 | 1093.8 | 599.2 |

| Gene ID | WT_HC vs *cbp1*-1_HC | | *cbp1*-1:*CBP1*_HC vs *cbp1*-1_HC | | WT_VLC 0.3 h vs *cbp1*-1_VLC 0.3 h | | *cbp1*-1:*CBP1*_VLC 0.3 h vs *cbp1*-1_VLC 0.3 h | | WT_VLC 2.0 h vs *cbp1*-1_VLC 2.0 h | | *cbp1*-1:*CBP1*_VLC 2.0 h vs *cbp1*-1_VLC 2.0 h | |
|---|---|---|---|---|---|---|---|---|---|---|---|---|
| | $log_2FC$ | FDR | $log_2FC$ | FDR | $log_2FC$ | FDR | $log_2FC$ | FDR | $log_2FC$ | FDR | $log_2FC$ | FDR |
| **Cre01.g047650** | 1.21 | 2.4.E-03 | 0.34 | 7.0.E-01 | 1.33 | 1.4.E-03 | 1.23 | 3.6.E-04 | 0.01 | 9.9.E-01 | 0.61 | 5.0.E-01 |
| **Cre03.g198250** | -1.68 | 7.2.E-09 | 0.27 | 7.3.E-01 | -2.61 | 2.9.E-13 | -0.50 | 1.1.E-01 | -3.09 | 9.7.E-16 | -1.35 | 6.0.E-04 |
| **Cre03.g204465** | 2.19 | 1.5.E-04 | -0.55 | 5.1.E-01 | 2.09 | 6.6.E-07 | 1.49 | 3.9.E-04 | 0.44 | 4.0.E-01 | 0.48 | 7.9.E-01 |
| **Cre03.g204577** | N.D. | N.D. | N.D. | N.D. | 2.34 | 8.0.E-07 | 2.56 | 9.3.E-09 | 0.21 | 8.0.E-01 | 1.57 | 1.9.E-01 |
| **Cre04.g216550** | 0.80 | 7.2.E-02 | 1.13 | 1.2.E-02 | 1.31 | 2.1.E-04 | 1.24 | 2.4.E-05 | 1.00 | 4.6.E-02 | 1.06 | 1.6.E-01 |
| **Cre04.g223250** | N.D. | N.D. | N.D. | N.D. | 2.44 | 3.7.E-09 | 2.69 | 2.1.E-17 | 1.88 | 2.8.E-05 | 1.37 | 7.1.E-02 |
| **Cre05.g245500** | 0.27 | 6.3.E-01 | -1.50 | 8.5.E-06 | -1.78 | 1.2.E-06 | -1.95 | 5.0.E-12 | -1.10 | 2.3.E-03 | -1.55 | 8.5.E-08 |
| **Cre08.g380700** | 1.91 | 2.5.E-09 | 0.79 | 8.4.E-03 | 1.90 | 5.5.E-06 | 1.25 | 6.4.E-04 | 0.52 | 4.9.E-01 | 0.76 | 3.2.E-01 |
| **Cre11.g469650** | 0.58 | 1.3.E-01 | -0.03 | 9.8.E-01 | 1.25 | 1.4.E-03 | 1.31 | 8.5.E-05 | 0.06 | 9.3.E-01 | 0.37 | 7.7.E-01 |
| **Cre12.g527250** | 0.82 | 8.7.E-02 | -0.85 | 1.1.E-01 | 1.85 | 2.2.E-05 | 1.60 | 1.2.E-05 | 1.31 | 3.1.E-03 | 0.76 | 1.9.E-01 |
| **Cre12.g555700** | N.D. | N.D. | N.D. | N.D. | 2.76 | 4.4.E-10 | 2.96 | 1.2.E-17 | -0.06 | 9.4.E-01 | 1.27 | 3.6.E-01 |
| **Cre16.g652200** | -1.99 | 3.6.E-06 | -0.86 | 6.0.E-02 | -2.96 | 2.1.E-15 | -1.13 | 2.1.E-04 | -1.46 | 9.6.E-03 | -0.72 | 3.1.E-01 |
| **Cre17.g719150** | 2.26 | 1.3.E-03 | -0.57 | 5.7.E-01 | -0.96 | 8.6.E-02 | -1.22 | 8.8.E-03 | -3.54 | 2.5.E-07 | -3.76 | 8.4.E-12 |
| **Cre19.g751047** | -0.93 | 1.6.E-03 | -0.34 | 6.0.E-01 | -2.17 | 1.2.E-09 | -1.14 | 1.0.E-05 | -2.92 | 8.7.E-12 | -2.05 | 2.7.E-06 |

Gene IDs represent the gene accession numbers from Phytozome.
